# Zinc Alters the Supramolecular Organization of Nucleic Acid Complexes with Full-Length TIA1

**DOI:** 10.1101/2023.01.25.525508

**Authors:** Yizhuo Yang, Keith J. Fritzsching, Sasha He, Ann E. McDermott

**Affiliations:** Department of Chemistry, Columbia University, New York, NY 10027, USA; Department of Organic Materials Science, Sandia National Laboratories, Albuquerque, NM 87185, USA

## Abstract

T-Cell Intracellular Antigen-1 (TIA1) is a 43 kDa multi-domain RNA-binding protein involved in stress granule formation during eukaryotic stress response, and has been implicated in neurodegenerative diseases including Welander distal myopathy and amyotrophic lateral sclerosis. TIA1 contains three RNA recognition motifs (RRMs), which are capable of binding nucleic acids and a C-terminal Q/N-rich prion-related domain (PRD) which has been variously described as intrinsically disordered or prion inducing and is believed to play a role in promoting liquid-liquid phase separation connected with the assembly of stress granule formation. Motivated by the fact that our prior work shows RRMs 2 and 3 are well-ordered in an oligomeric full-length form, while RRM1 and the PRD appear to phase separate, the present work addresses whether the oligomeric form is functional and competent for binding, and probes the consequences of nucleic acid binding for oligomerization and protein conformation change. New SSNMR data show that ssDNA binds to full-length oligomeric TIA1 primarily at the RRM2 domain, but also weakly at the RRM3 domain, and Zn^2+^ binds primarily to RRM3. Binding of Zn^2+^ and DNA was reversible for the full-length wild type oligomeric form, and did not lead to formation of amyloid fibrils, despite the presence of the C-terminal prion-related domain. While TIA1:DNA complexes appear as long “daisy chained” structures, the addition of Zn^2+^ caused the structures to collapse. We surmise that this points to a regulatory role for Zn^2+^. By occupying various “half” binding sites on RRM3 Zn^2+^ may shift the nucleic acid binding off RRM3 and onto RRM2. More importantly, the use of different half sites on different monomers may introduce a mesh of crosslinks in the supramolecular complex rendering it compact and markedly reducing the access to the nucleic acids (including transcripts) from solution.

## INTRODUCTION

The T-Cell Intracellular Antigen-1 (TIA1) protein is pivotal for response to stress and stress granule formation in eukaryotic cells. It has three RNA recognition motifs (RRM) which are capable of binding nucleic acids and a Q/N-rich low complexity prion-related domain (PRD) (1,2) which has been described as intrinsically disordered or prion inducing and confers liquid-liquid phase separation tendency. connected with the assembly of stress granule formation. Mutations in TIA1 and has been implicated in neurodegenerative diseases including Welander distal myopathy and amyotrophic lateral sclerosis.

Stress granules (SGs) are membrane-less reversibly formed biomolecular condensates of RNA-binding proteins and translationally stalled mRNAs.(3,4) They are observed in the cytoplasm when the cell responds to stress conditions, such as heat shock, osmotic stress, or oxidative stress, and TIA1 is a key protein component. After stress is relieved, stress granules disassemble allowing mRNA translational repression to be reversed.(5) The process of SG formation and dissolution affects a range of biological activities and helps cells overcome environmental challenges. Disease related mutations the key RNA-binding proteins of stress granules including TIA1 cause the stress granules to become less dynamic and soluble, and become prone to amyloid nucleation in neurodegenerative diseases.(6-8)

Increasing evidence suggests that liquid-liquid phase separation of TIA and other proteins underlies the formation of stress granules, analogous to many other membrane-less cellular structures.(9) Though the complex network of molecular interactions among their components makes it hard to determine precise mechanistic details of SGs formation, models have been proposed for SGs assembly, and the low-complexity domains of the RNA-binding proteins such as TIA1 appear to be key regulators of liquid-liquid phase separation.(10,11) The low-complexity regions are typically enriched in a limited number of amino acid types, primarily glycine (G), glutamine (Q), asparagine (N), tyrosine (Y), and serine (S). Phase separation of RNA-binding proteins including TDP43(12), FUS(13), is mainly affected by multivalent interactions among aromatic residues (typically Tyrosine) from prion-like domains and positively charged residues (typically Arginine) from RNA-binding domains.(14,15) A regulatory role for the low complexity domain is consistent with the observations that isolated prion-like domains and isolated RNA-binding domains both have very different phase behavior than the full-length proteins. Apart from the interactions mentioned above, RNA-binding domains are also proposed to engage in a variety of other interactions that promote assembly of SGs, including interactions with nucleic acids and with other RNA-binding proteins. Also, post-translational modifications at sites in the RNA-binding domains can contribute to the regulation of SGs.(16)

Though TIA1 is considered to be vital for stress granule formation, its full-length structure is not known. Two of the three RRMs have been studied structurally as excised domains while the Q/N-rich low complexity prion-related domain (PRD) and the first RRM have been more elusive.(1,2) The RRM domains are relatively stable globular domains separated by linkers and responsible for interacting with RNA. The PRD domain is believed to be an intrinsically disordered region and plays an important role in liquid-liquid phase separation that leads to the assembly of SGs.(17) Biophysical techniques indicate that full-length TIA1 proteins form oligomers under many conditions both in vivo and in vitro, for example when binding with nucleic acids and Zn^2+^. The three RRM domains of have different binding affinities and specificity towards oligonucleotides in this and other systems.(18) For TIA1, it is believed that RRM2 plays a dominant role in interacting with U-rich RNAs, while RRM3 recognizes C-rich RNAs.(19) The functional contributions of RRM1 and the PRD remain unclear. One hypothesis is that RRM1 has no tendency to bind with RNA due to the presence of degenerate RNP motifs in the sequence, with either modifications in normally conserved critical amino acids, and/or reduced thermostability.(1) The lower conservation of RRM1 compared with RRM2 and RRM3 makes it more likely that it serves other functions for example contributing to recruiting other proteins into stress granules by protein-protein interactions, rather than interact with RNA directly.(20) Previous solution-state NMR studies on a TIA1-RRM1,2,3 construct upon binding with U-rich RNAs revealed that RRM2 and RRM3 cooperate in binding to RNA ligands.(1) Residues in the noncanonical N-terminal helix α_0_ of RRM3 experience strong chemical shift perturbations upon RNA binding in the context of TIA1-RRM2,3 construct, while the isolated TIA1-RRM3 construct doesn’t exhibit similar strong chemical shift perturbations for α_0_ upon RNA binding, which indicates that the role of α_0_ in RNA recognition and the RNA binding process depends on cooperative interactions involving both RRM domains. RNA binding is partly controlled by pH changes due to the protonation/deprotonation of RRM3 histidine residues. The histidine residues are highly conserved among TIA1 homologs, especially H259. The protonation/deprotonation of H259 can disrupt the cation-π contacts between H259 and W283, thus affecting RNA recognitions.(21)

The studies above illustrate the importance of studying nucleic acid binding in the full-length TIA1 protein (rather than in excised domains). Notably, most phase separating proteins have multiple binding sites. Interactions between the RRM domains as well as interactions between the RRMs and the low complexity domain strongly modulate function. Intermolecular interactions are also important since TIA1 forms oligomers and phase separates under many bio-relevant conditions. These multiprotein interactions are expected to have functional consequences. For example, oligonucleotide binding to TIA1 smaller constructs with only a RRM single binding site induce small conformational changes, and apparently do not enhance phase separation.(22) By contrast, constructs with multiple binding sites have been seen to contribute to phase separation for TIA1 and other proteins. In other words, oligonucleotide binding site stoichiometry and multivalency, rather than the length of the oligonucleotides or specific binding motifs might be critical for the connection between binding and phase separation.(22) Therefore, the study of structure and function of the domains in context of full-length oligomeric constructs is expected to be important for understanding TIA1.

The fact that Zn^2+^ concentrations are highly elevated in neurodegenerative brains and in stress conditions (such as arsenite) makes Zn^2+^ also an interesting target for studies of regulatory ligand binding. Ambient Zn^2+^ influences the oligomerization rate of TIA1 and the size of oligomers, independent of changes in β sheet content or the presence of RNA.(23) Self-multimerization of TIA1 induced by Zn^2+^ can also be reversed by a Zn^2+^ chelator, such as N,N,N’,N’-tetrakis-(2-pyridylmethyl)ethylenediamine (TPEN). Addition of TPEN after Zn^2+^ treatment causes collapse of TIA1 oligomers.

Although both oligonucleotides and Zn^2+^ have been identified as important regulators of TIA1 oligomerization and stress granule formation, it is not clear that how they influence each other in the biomolecular condensates. While several excised TIA1-RRM constructs have been characterized with small-angle scattering and solution-state NMR to great benefit (24), to date there have not been structurally detailed studies of full-length TIA1 oligomers binding to oligonucleotides or Zn^2+^.

In our previous work using multidimensional solid-state NMR methods, we characterized full-length (fl) TIA for the first time. We showed that fl TIA1 protein forms a dense colloidal-like phase, that we described as micelle-like.(25) The micelle-like phase was formed on the timescale of hours when induced by concentration and agitation, and was stable for days, and its formation could be reversed to some extent by dilution. By contrast, in our hands the fl TIA1 did not form amyloids under mild buffer conditions. We observed that the two DNA binding modules, RRM2 and 3 were folded and solvent exposed in micellar fl TIA1. Therefore, in this study we examined the properties of the micellar form of fl TIA1 in more detail to test for competent ligand binding of single strand DNA and Zn^2+^, and the consequences of DNA and Zn^2+^ binding for oligomerization and conformational change.

## MATERIAL AND METHODS

### Protein Preparation

TIA1 proteins were expressed in the BL21DE3 bacterial cell line from the pET28b plasmid (Addgene) containing the Mus musculus (mouse) TIA1 sequence and an N-terminal His_6_-SUMO tag. For preparing uniformly ^13^C and ^15^N enriched samples, the cells were grown in 4 × 1L LB culture (25g/L LB, 50 μg/mL kanamycin sulfate) at 37°C with aeration by shaking at 255 RPM until the OD at 600 reached 0.65. Then the cells were harvested by centrifugation (3600 RPM, 25 min., 4°C), and gently resuspended into 1L of isotopically enriched M9 buffer (13.6g sodium phosphate, 6g potassium phosphate, 1g NaCl, 100μL 1M CaCl_2_ stock, 2mL 1M MgSO_4_ stock, 20mL of Solution C, 10mL 100x MEM Vitamin Solution (Gibco™), 50 μg/mL kanamycin (Thermo Scientific™), 1g ^15^N-NH_4_Cl (Cambridge Isotope Laboratories, Inc.), and 3g U-^13^C-D-Glucose (Cambridge Isotope Laboratories, Inc.), pH 7.4). After incubating at 37°C with shaking at 255 RPM for 45 min, 1mM IPTG (Thermo Scientific™) was added to induce protein expression. The cells were harvested again by centrifugation after expression for 4 hours. 40mL lysis/wash buffer (50mM sodium phosphate, 500mM NaCl, 20mM imidazole, 0.002% sodium azide, 1mM DTT, pH 7.4) was used to resuspend the cell pellets. The pellets were stored at -80°C overnight. The cells were defrosted and then lysed with 1mg/mL Lysozyme (Sigma-Aldrich), 1 tablet of EDTA-free protease inhibitors (Thermo Scientific™), and 2U of DNase (Thermo Scientific™). After reacting at room temperature for 30 minutes, the cell suspension was poured into the cylinder of the Emulsiflex C3 high pressure homogenizer and cycled in the system at 4 °C with the use of the connected cooling system. The homogenizing pressure was slowly increased to 40 psi during the process. The homogenized cell lysate was collected after ten minutes (during which 4 passes through the homogenizer were performed) and separated by ultracentrifugation (Type 70 Ti Fixed-Angle Titanium Rotor, 32,000 RPM (105,513 g), 45 min., 4°C). The supernatant containing the His_6_-SUMO-TIA1 was sterile filtered (0.2 μM) (Thermo Scientific™) before purification using Fast protein liquid chromatography (FPLC) (Bio-Rad Laboratories, Inc.). The supernatant was loaded onto a 1mL HisTrap nickel-affinity column (Cytiva Lifescience™) at 1mL/min, washed with more than 10 column volumes of lysis/wash buffer and further washed with a linear gradient up to 25% elution buffer (50mM sodium phosphate, 500mM NaCl, 300mM imidazole, 0.002% sodium azide, 1mM DTT, pH 7.4). The protein was eluted with 100% elution buffer as the final step. After elution, the His_6_-SUMO-TIA1 protein was mixed with 2U of SUMO protease (gift from New York Structural Biology Center) in order to cleave the His_6_-SUMO tag. The mixture was dialyzed in a 20,000 MW cut off cassette (Thermo Scientific™) for three hours in 1L exchange buffer (50mM sodium phosphate, 1M NaCl, 0.002% sodium azide, 0.5mM EDTA, 1mM DTT, 10% glycerol, pH 7.4) at room temperature, and transferred to another 1L exchange buffer for overnight dialysis at 4°C. The tag and cleaved proteins were then separated using nickel-affinity chromatography by repeating the FPLC method described above, except that the buffer used at the wash steps was exchange buffer and the buffer used at the elution step was exchange buffer with 300mM imidazole (Thermo Scientific™). Purified TIA1 protein was collected in the wash steps. Finally, the buffer was exchanged for the NMR buffer (25mM PIPES, 50mM NaCl, 0.5mM EDTA, 5mM TCEP, 0.002% sodium azide, 20% glycerol, pH 6.8) using a desalting column (HiPrep 26/10 desalting column from Cytiva Lifescience™). The protein was either used immediately or rapidly frozen with liquid nitrogen and stored at -80°C.

### Optical density assay

Assessment of the extent of protein oligomerization was based on monitoring absorbance at 550 nm using BioTek Plate Reader (Agilent BioTek NEO2) to reflect turbidity changes induced by protein oligomer formation. Full-length TIA1 at a concentration of 15 uM was prepared in NMR buffer without EDTA. 80 uL protein was added into each well of a 96-well plate. To study the rates and amount of oligomerization at different conditions, DNA, ZnCl_2_, and NMR Buffer were added to various wells. The total volume of the mixture in each well was same (100 uL). The absorbance at 550nm was recorded every 40 seconds for 3 hours at room temperature while shaking (using the Double Orbital Continuous shaker in the BioTek Plate Reader, with a frequency of 282 cpm and amplitude of 3 mm).

### Isothermal titration calorimetry

Isothermal titration calorimetry (ITC) was applied to characterize binding stoichiometry and affinity of TIA1 towards DNA. All titrations were carried out using MicroCal Auto iTC200 calorimeters (Malvern Panalytical). To provide a consistent environment, TIA1 protein were in the same buffer used for NMR structural study (NMR buffer). 10uM protein was titrated with 200uM ssDNA at 298K. The same buffer was used for protein and ligands to minimize ΔH contributions from dilution. Raw data was analyzed to produce a plot of ΔH per mole of injection versus molar ratio using the Origin ITC Analysis software. Data were fit as either a one-site or two-site binding model to provide information of binding affinity thermodynamics and stoichiometry. Sigmoidal curves were typically observed during the titrations.

### Negative stained electron microscopy

Transmission Electron Microscopy was used to identify the morphology of oligomeric fl TIA1 under different conditions. Two-fold diluted solutions of oligomeric TIA1 sample were applied to carbon-coated copper grids and stained using 2% uranyl acetate. The residual stain was removed by back-blotting. Electron microscopy images of each grid were collected using a JEOL JEM-1230 microscope operating at 120kV equipped with a Gatan US-4000 charge-coupled device detector.

### Solid-state NMR

All solid-state NMR experiments were carried out using a 750 MHz Bruker NEO NMR spectrometer with a 1.9 mm Bruker HCN probe. The variable temperature set point was 235K, and the sample spinning frequency was typically 16.666 kHz. Under these conditions the sample temperature was estimated to be 265K due to frictional heating.(26) After optimizing the parameters of the pulses, ^1^H 90-pulse was 2.3 μs at 53W, the ^13^C 90-pulse was 4.5 μs at 80W, and the ^15^N 90-pulse was 4.1 μs at 120W. Cross polarization (CP) conditions were optimized with tangential ramped shaped pulses, contact times of 1200 μs for ^13^C[^1^H] and 900 μs for ^15^N[^1^H] were used. For NCA transfer, specific CP was optimized near the ν_1_(^13^C) =1.25 ν_r_ and ν_1_(15N) = 0.25 ν_r_ matching condition, with a 3.5 ms tangential ramp on the carbon channel. Swept-frequency two-pulse phase modulation (SWFTPPM) decoupling with ν_1_(^1^H) = 100 kHz was used during acquisition. The direct carbon dimensions were acquired for 13 ms. In 2D ^13^C-^13^C DARR experiments, the final indirect dimension acquisition time was 6.75 ms. In 2D NCA experiment the final indirect dimension acquisition time was 11.5 ms. NMR data were processed using TopSpin (Bruker) and analyzed using CCPN.(27)

## RESULTS

### Binding of full-length TIA1 to a TC-rich single strand DNA

To characterize DNA binding to TIA1, we designed a single stranded DNA oligomer that is expected to be competent for binding both RRM2 and RRM3, namely: 5’-TTTTTACTCC-3’ (which we refer to as TC1, containing one TC-rich binding site). RRM2 and RRM3 were previously shown to bind to U-rich RNAs and C-rich RNA motifs (such as 5’-ACUCC-3’) respectively using surface plasmon resonance.(24) DNA is less susceptible to degradation than RNA due to common exposure to RNase contamination from environmental sources during sample preparation and characterization. DNA rather than RNA was used here as they have comparable binding affinities but DNA is better suited to the long experimental times characteristic for SSNMR.

Isothermal titration calorimetry (ITC) was used to characterize binding stoichiometry and affinity of TIA1 towards ssDNA. A sample containing 10μM fl TIA1 was titrated with 200μM TC1 at 298K. A sigmoidal titration curve (Fig. 1) was typically observed for protein-ligand binding when no oligomerization formed. Comparing different fitting models, we concluded that two-site binding models fit somewhat better than a one-site fitting models with respect to <χ>^2^ criteria. Thermodynamic parameters obtained from the fitting models are shown in table 1 and 2. The overall negative binding enthalpy indicates that the DNA binding is enthalpically driven under these conditions. The dissociation constant of K_D2_ =116 nM suggests that TC1 binds to TIA1 with a relatively strong binding affinity, while the value of N =1 indicates a 1:1 binding stoichiometry for that binding mode. The value of K_D1_, 7.4 mM, is associated with less well-defined features in the curve, and is less confidently fit and less consistent over multiple data sets. Overall, the ITC experiments confirm that TC1 binds to TIA1 with strong binding affinity and a stoichiometry of 1 to 1, with additional weaker binding interactions that are less well characterized by these experiments. Binding affinities for Zn^2+^ to TIA1 were not successfully measured with ITC because this ligand induced oligomerization during the titration. The heats released from oligomerization likely dominated observed heats released upon binding, making the contribution from the metal interactions difficult to isolate. More importantly the oligomerization is strongly dependent of agitation, and practically difficult to record because of instrument clogging and other technical challenges.

**Table 1.** Thermodynamic values for a one-site fitting model of TC1 bound to TIA1

| K (M<sup>-1</sup>) | K<sub>D</sub> (nM) | ΔH (kcal/mol) | N |
| --- | --- | --- | --- |
| $3.79 \cdot 10^6 \pm 8.79 \cdot 10^5$ | $263 \pm 60$ | $-30.7 \pm 0.8$ | $1.07 \pm 0.02$ |

**Table 2.** Thermodynamic values for a two-site fitting model of TC1 bound to TIA1

| $K_1$ ($M^{-1}$) | $K_{D1}$ (mM) | $\Delta H_1$ (kcal/mol) | $K_2$ ($M^{-1}$) | $K_{D2}$ (nM) | $\Delta H_2$ (kcal/mol) | $N_2$ |
| --- | --- | --- | --- | --- | --- | --- |
| $134 \pm 64300$ | $7.46 \pm 3 \cdot 10^3$ | $-659 \pm 129 \cdot 10^5$ | $8.62 \cdot 10^6 \pm 1.04 \cdot 10^9$ | $116 \pm 1 \cdot 10^4$ | $-29.9 \pm 2.5$ | $1.03 \pm 0.15$ |

**Figure 1.**
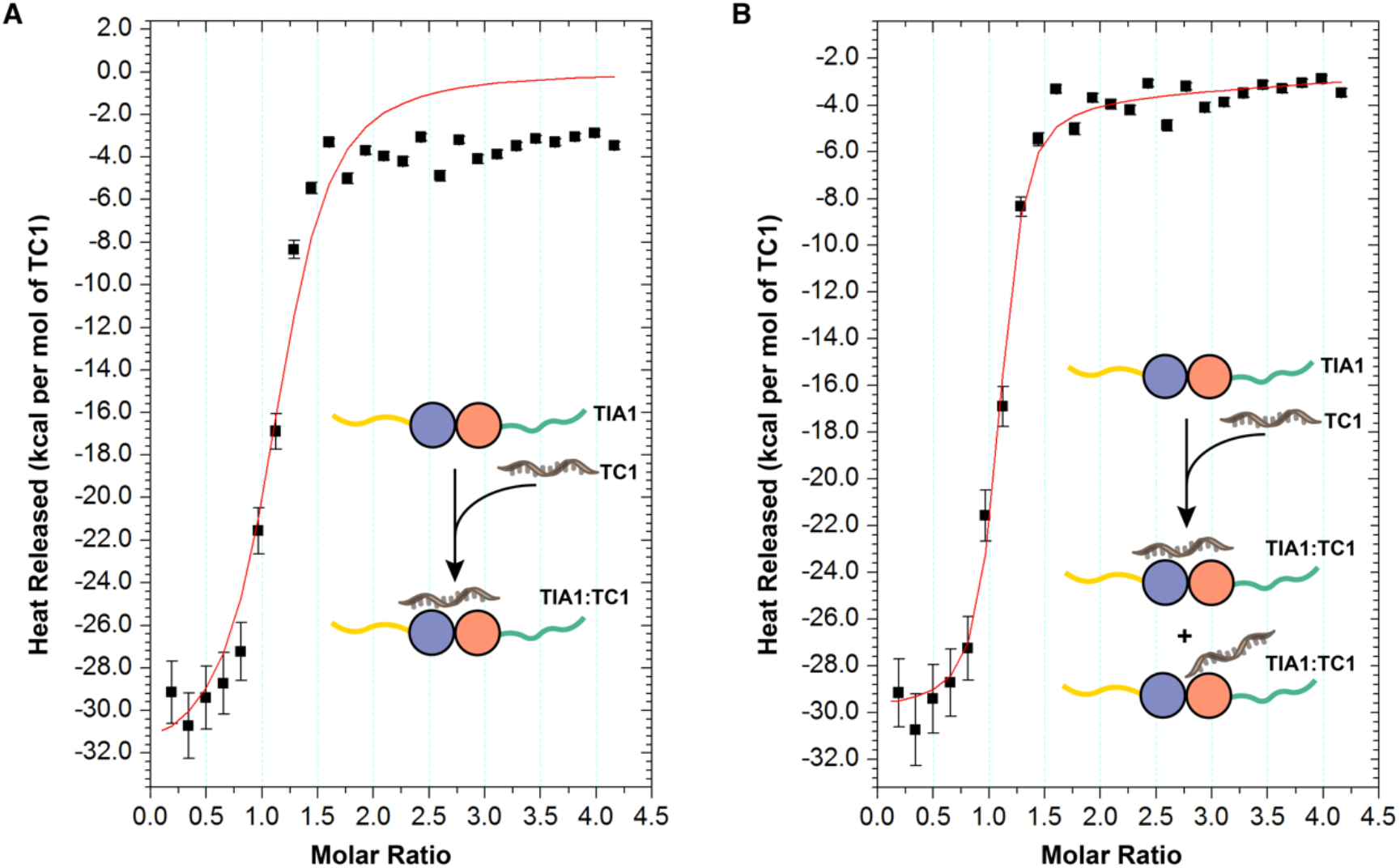
Representative binding isotherms and binding-model fits for titration of TIA1 with ssDNA TC1. (A) One site fitting model of TC1 bound to TIA1. (B) Two sites fitting model of TC1 bound to TIA1.

### Oligomerization assays

We also asked whether the presence of Zn^2+^ or TC1 DNA promotes oligomerization or phase separation of TIA1. We monitored oligomerization in a qualitative fashion by recording optical turbidity (A550 nm) in a UV/VIS optical spectrophotometer.(28) Zn^2+^ was added using 1.4 mM ZnCl_2_ stock solution to a final concentration of 350 μM. The buffer contained 50 mM Cl^-^, so the Cl^-^ concentration was only changed by a marginal amount and would not be expected to affect protein behavior. Increasing the protein concentration from 0 to 8 μM accelerated the oligomerization of TIA1 when Zn^2+^ concentration in the buffer was held constant (and was in a molar excess relative to the protein) (Fig. S1). Maximum turbidity (A550 nm) of TIA1 was directly proportional to the concentration of TIA1 (Fig. 2A). We interpret this result to mean the size of the oligomers didn’t change ostensibly when the protein concentration increased. When the protein concentration was higher (> 10 μM), Zn^2+^-induced oligomers formed immediately even before the start of recording and reached the maximum turbidity within seconds, leading to an inaccuracy in recording the maximum turbidity. As the concentration increases above 10 μM, the maximum turbidity is difficult to measure accurately because of the sample settling gravitationally. When the protein concentration was held constant, increasing the Zn^2+^ concentration promoted TIA1 oligomerization dramatically (Fig. 2B). The log of maximum turbidity of TIA1 exhibited proportionality to the concentration of Zn^2+^, as expected in simple binding curves, consistent with the size of the oligomers becoming larger as more Zn^2+^ was added. In figure 2C, excess Zn^2+^ and 10nt TC1 ssDNA were added to TIA1 in separate samples, and in a separate experiment were added simultaneously to induce oligomerization of TIA1. The final composition of the mixture was 2 μM TIA1, 95 μM Zn^2+^, and 15 μM TC1 ssDNA. While TC1 alone did not enhance the oligomerization of TIA1, addition of Zn^2+^ alone quickly caused the formation of TIA1 oligomerization (within 5 min) with the absorbance at 550 nm increasing rapidly. We conclude that ssDNA with only one TC rich element does not induce oligomerization of TIA1. On the other hand, with agitation, TC1 bound to TIA1 caused notable oligomerization with a rising turbidity. Furthermore, when TC1 was added into Zn^2+^-induced oligomers TIA1, the mixture became less opaque, and the absorbance at 550 nm decreased significantly (Fig. 2D). When the order of addition was reversed, which is to say Zn^2+^ was added to a TIA1:TC1 complex, the turbidity showed an increase. Both protocols, regardless of the order of addition, resulted in similar final complexes as assessed by the turbidity, suggesting reversible effects on organization.

**Figure 2.**
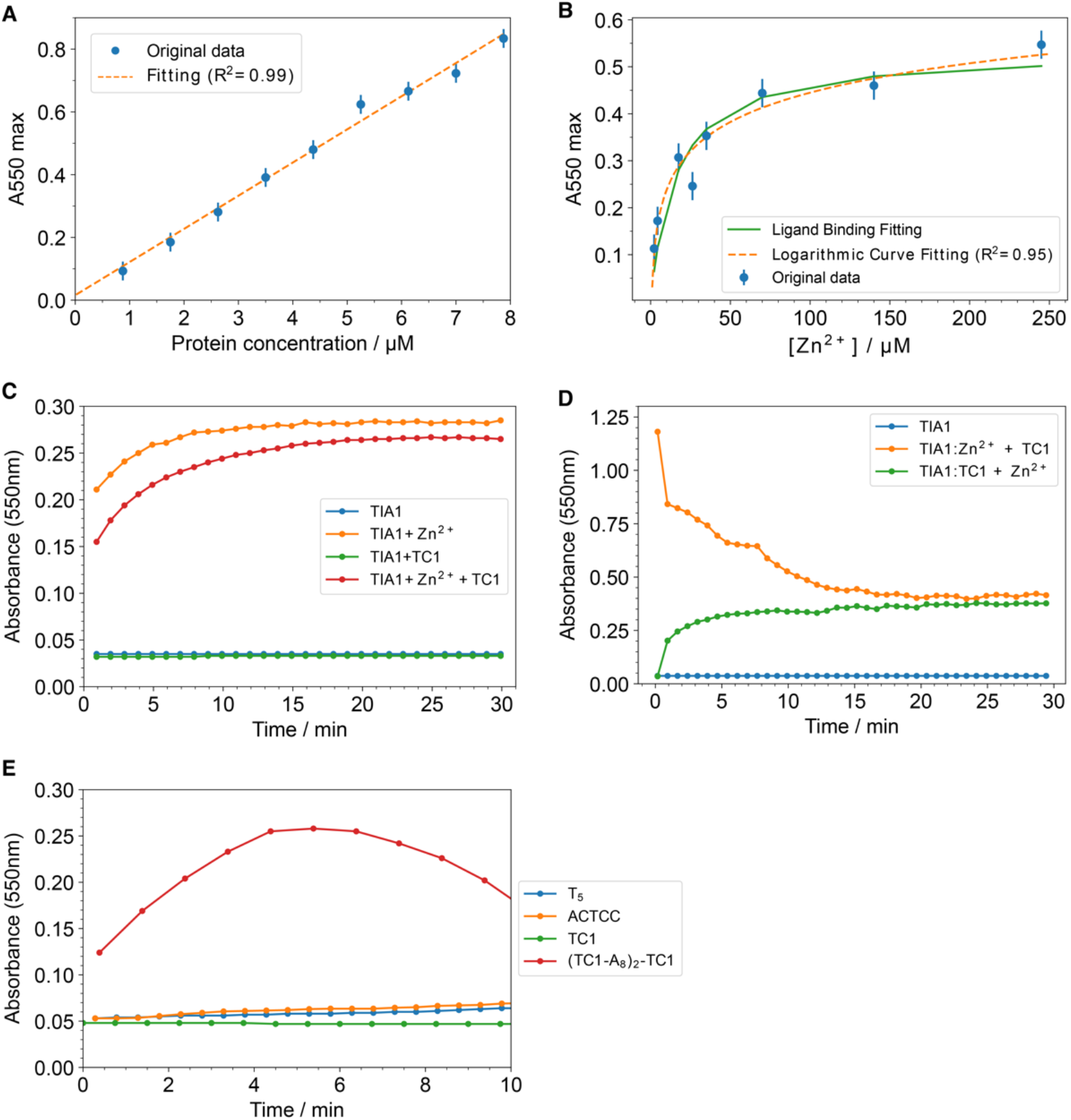
Oligomerization assays of TIA1 under different conditions. (A) Maximum turbidity (A550 nm) of 0-8 μM TIA1 with excess ZnCl_2_. The exact concentrations of TIA1 in each well are shown in table S1. The concentration of ZnCl_2_ was 350 μM. The buffer contained 25mM PIPES, 50mM NaCl, 5mM TCEP, 0.002% sodium azide, 20% glycerol, pH 6.8. The turbidity of buffer with no protein was measured for reference. (B) Maximum turbidity (A550 nm) of 5 μM TIA1 with different ZnCl_2_ concentrations. The exact concentrations of Zn^2+^ in each well are shown in table S2. The data can be fit to a logarithmic curve (*y* = 0.09 In *x* + 0.03) with R^2^=0.95. The binding data can also be fit to a simple binding model (*y* = (*B*_*max*_ × *x*)/(*K*_*d*_ + *x*)) with K_d_ =15.9 ± 5.3 μM, where B_max_ is the maximum specific binding in the same units as y and K_d_ is the equilibrium dissociation constant in the same units as x. Our reaction is probably more complicated in that the binding induces a change in oligomerization (based on turbidity), for example: m(TIA1)_n_ + m*nZn^2+^ = [TIA1: Zn^2+^]_m*n_. (C) Turbidity of TIA1, TIA1 with ZnCl_2_, TIA1 with TC1 ssDNA, and TIA1 with ZnCl_2_ + TC1 ssDNA. The final concentration of each component was 2 μM TIA1, 100 μM Zn^2+^, and 15 μM TC1 ssDNA. Zn^2+^ alone promoted the increase of turbidity faster and higher than both Zn^2+^ and ssDNA. 10nt TC1 ssDNA alone did not induce changes of turbidity. (D) Turbidity of TIA1 with ligands adding in a different order. The measurement was performed with a final concentration of TIA1 at 12 μM, Zn^2+^ at 100 μM and TC1 ssDNA at 24 μM. The addition of TC1 to the mixture of TIA1 and Zn^2+^ caused a rapid decrease of turbidity (orange curve). On the other hand, the addition of Zn^2+^ to the mixture of TIA1 and TC1 caused a moderate increase of turbidity immediately (green curve). Both of them formed a final complex with the turbidity at around 0.4, indicating that similar size of oligomers formed at the equilibration state no matter what order TC1 and Zn^2+^ were added to TIA1. (E) Turbidity of TIA1 with different ssDNA sequences. The DNA sequences used in the oligomerization assay and the maximum turbidity (A550 nm) of each TIA1:DNA complex were listed in table 3. Both protein and ssDNA concentration were at 20 μM. ssDNA with multiple binding sites caused more oligomerization regardless of the length of the nucleotides.

We explored sequence preferences in nucleic acid binding in the absence of Zn^2+^. A series of ssDNA constructs with single or multiple binding sites were compared in terms of binding and oligomerization (Table 3, Fig. 2E and Fig. S2). ssDNA with a length of 10nt was shown in previous work to bind well and exhibits clear composition preferences. DNA sequences with multiple Ts are expected to preferentially bind RRM2 while those with multiple Cs are targeted to RRM3.(19) Accordingly, T_5_, T_5_A_5_ and T_10_ (see Table 3) were designed to bind to RRM2; ACTCC, C_5_, A_5_ACTCC and C_10_ was designed to bind to RRM3. Other constructs were designed to bind to multiple sites, having multiple binding elements connected with linkers. DNA sequences with a length of 5nt (which we expect might be sufficient in length for binding to a single binding site but would be insufficient in length for spanning two binding sites) did not contribute to oligomerization of TIA1. T_10_-A_2_-C_10_ and C_10_-A_2_-T_10_ caused the most effective and rapid oligomerization based on the turbidity assay (Fig. S2); presumably this is because these sequences bind to both RRM2 and RRM3 with high affinity and link several monomers together. DNA sequences with three binding sites and separated with long A linker ((TC1-A_8_)_2_-TC1) also promoted oligomerization effectively (Fig. 2E). It should be noted that following oligomerization, in the absence of stirring, the turbidity gradually dropped on a minutes-to-hours timescale as the oligomers settle down to the bottom by gravity. DNA T_10_, which is expected to bind to RRM2, induced oligomerization of TIA1 as indicated by increased turbidity, while DNA C_10_ which is targeted for RRM3 did not appreciably affect the turbidity (Fig. S2). Therefore, possibly RRM2 plays a more essential role in DNA binding and inter-monomer linking than RRM3 in the absence of Zn^2+^.

**Table 3.**
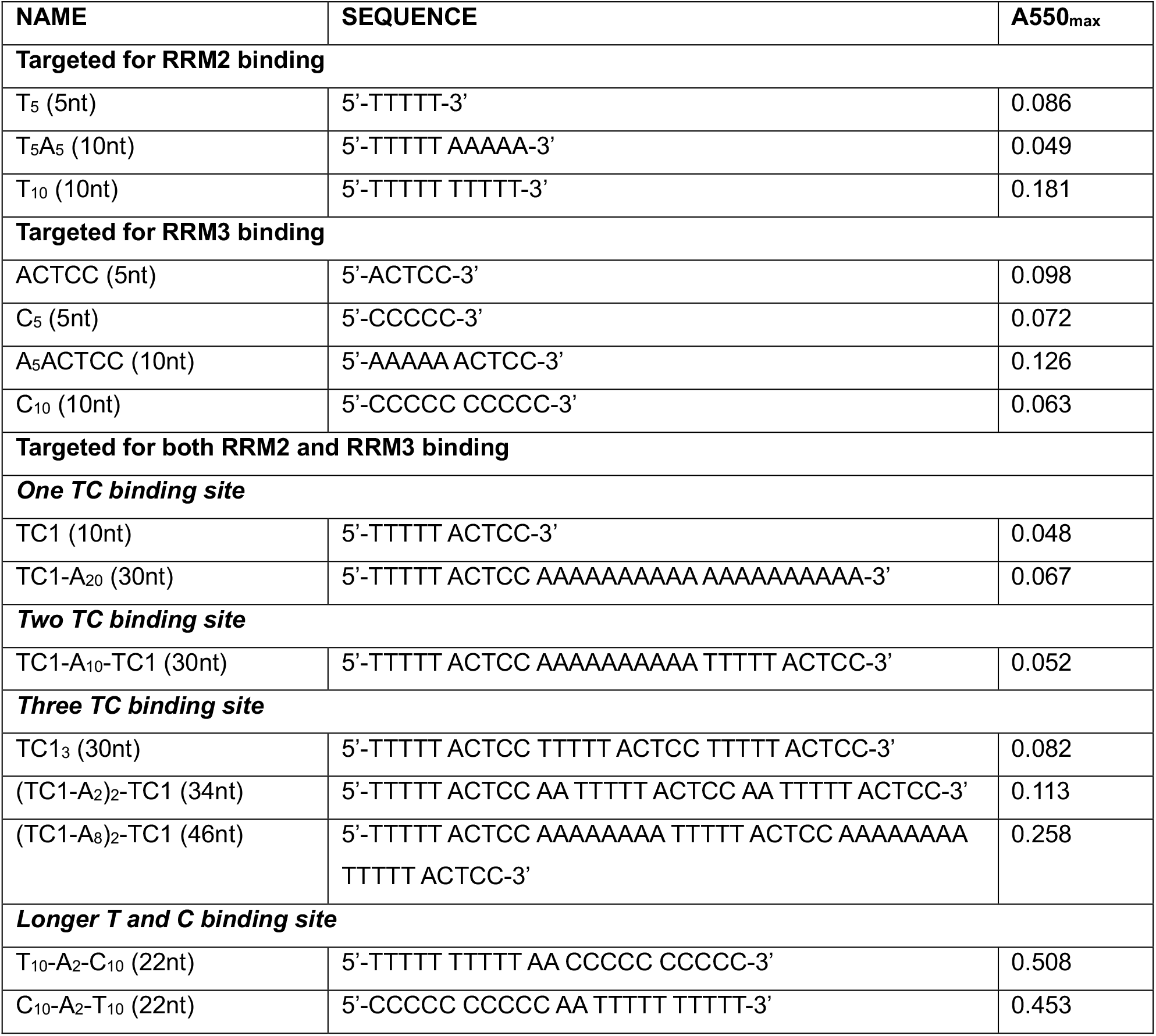
Sequences of ssDNA used in the oligomerization assay

### Identification the TIA1 oligomerization morphology with Transmission Electron Microscopy

Transmission electron microscopy (TEM) was used to image TIA1 oligomers, including those prepared under essentially identical conditions to the samples for the oligomerization assays and solid-state NMR experiments. All samples described were prepared with stirring with a magnetic stirring bar for two days at 4°C to induce oligomerization before blotting and staining. The protein oligomers were negatively stained on carbon-coated copper grids and screened with a JEOL JEM-1230 microscope. Electron Microscopy images of fl TIA1 oligomeric samples show no evidence of formation of amyloid fibrils in the presence nor the absence of TC1 or Zn^2+^ (or both). In figure 3, Panel A shows images of oligomeric apo TIA1. The typical apparent dimension of apo TIA1 micelles or particles in panel A is 20 nm (measured with software processing). Panel B shows images of TC1 bound TIA1 oligomers. With TC1 bound to TIA1, the oligomers appear linked, forming strings (>200 nm) with DNA presumably acting as a scaffold. Panel C shows oligomeric Zn^2+^ bound forms of TIA1. When TIA1 oligomerizes in the presence of Zn^2+^, oligomers form large irregular droplets. The average dimension of Zn^2+^ bound forms of TIA1 in C is 50 nm, with each droplet separated from the others. It corresponds with the result from the oligomerization assay that with the addition of Zn^2+^, in that turbidity of TIA1 increases dramatically and the oligomers formed apparently tended to settle down quickly due to gravity. Panel D shows the oligomers of fl TIA1 bound to both TC1 and Zn^2+^. When oligomerized with both TC1 and Zn^2+^, TIA1 forms round particles of apparent typical diameter 37 nm, larger than for apo TIA1 oligomers, but smaller than for Zn^2+^ bound forms of TIA1 oligomers. The relative particle sizes appear to explain the trends in changes in turbidity: TIA:TC1 (with agitation) >> TIA1:Zn^2+^ (50 nm) > TIA1:TC1:Zn^2+^ (37 nm) > TIA:TC1 (without agitation) ∼ apo TIA1 (20 nm).

**Figure 3.**
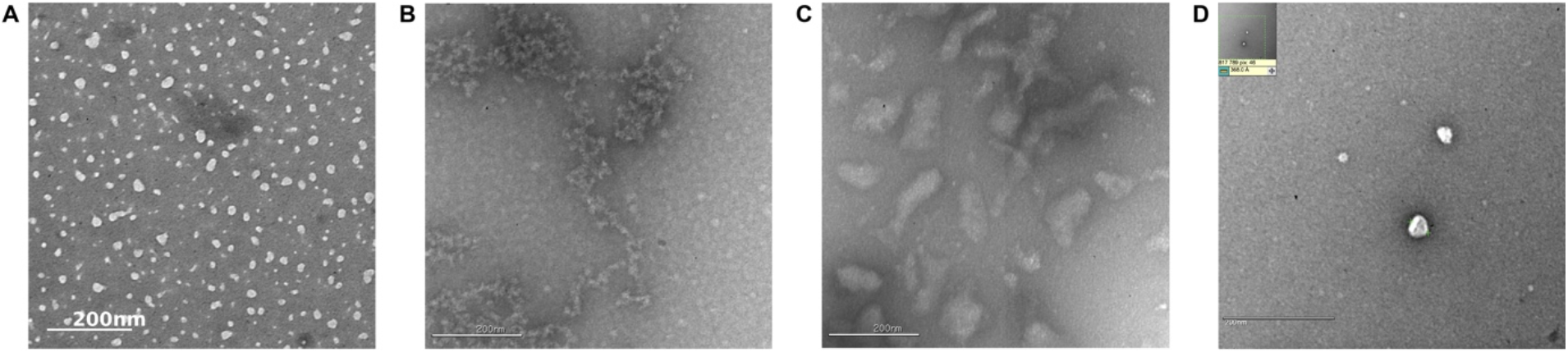
(A-D) Negatively stained electron micrographs of (A) TIA1, (B) TIA1 with TC1 /EDTA, (C) TIA1 with ZnCl_2_ /no EDTA, and (D) TIA1 with TC1 and ZnCl_2_ /no EDTA.

### Comparison of TC1 bound and apo forms of TIA1 by SSNMR

Solid state NMR experiments allowed us to further investigate the role of each RRM in DNA binding in the context of the oligomeric state. NMR allows information on the binding in an atom-by-atom, residue-by-residue fashion. For the NMR experiments, a twofold excess TC1 was mixed with TIA1. Although 10nt TC1 ssDNA did not particularly promote the oligomerization of TIA1, we allowed the oligomeric assemblies to slowly and reversibly settle to form a gel like state as we had done previously for the apo oligomeric system. The settled sample (soft pellet) was packed into the NMR rotor by ultracentrifugation and the supernatant was removed. Presumably, excess TC1 that did not bind would have remained in the supernatant and was not included.

In figure 4 we display the 2D ^13^C-^13^C correlation spectra of fl apo TIA1 (red) and of fl TIA1 with 10nt TC1 ssDNA (black). Labels highlight peaks that differ between the apo and TC1 bound forms. Numerous peaks appear only in the presence of the DNA. Side chain crosspeaks (Cα-Cγ, Ca-Cβ, Cβ-Cγ) of I117, I122, P128, V138, I164, I177 in RRM2, and V205, P238, I243, V269, V279 in RRM3 appear in the presence of TC1, but not in the absence, and Cα-Cβ crosspeaks for T118 and T179 in RRM2 increase in intensity due to addition of TC1. Another peak that appears in the spectrum due to TC1 binding is tentatively assigned as Cα-Cβ (Cα 58.65 ppm and Cβ 72.43 ppm) for T19 in RRM1, on the basis of the agreement with predicted shifts (Cα 59.25 ppm and Cβ 72.19 ppm). In general, for dipolar based spectra such as this CP DARR spectrum(29) appearance of a peak would suggest that the system has become more ordered or static (and exhibits reduced dynamics), while disappearance would suggest that dynamics have been increased. A shift in the residue’s position suggests that the ligand binds in close contact with the residue, or that allosteric effects cause changes in the residues environment(30). In the following analysis we interpret any of these effects on the crosspeak as an indication of binding, and as indicative of the residue’s likely involvement in the binding surface, or a related allosteric change in protein structure and dynamics.

**Figure 4.**
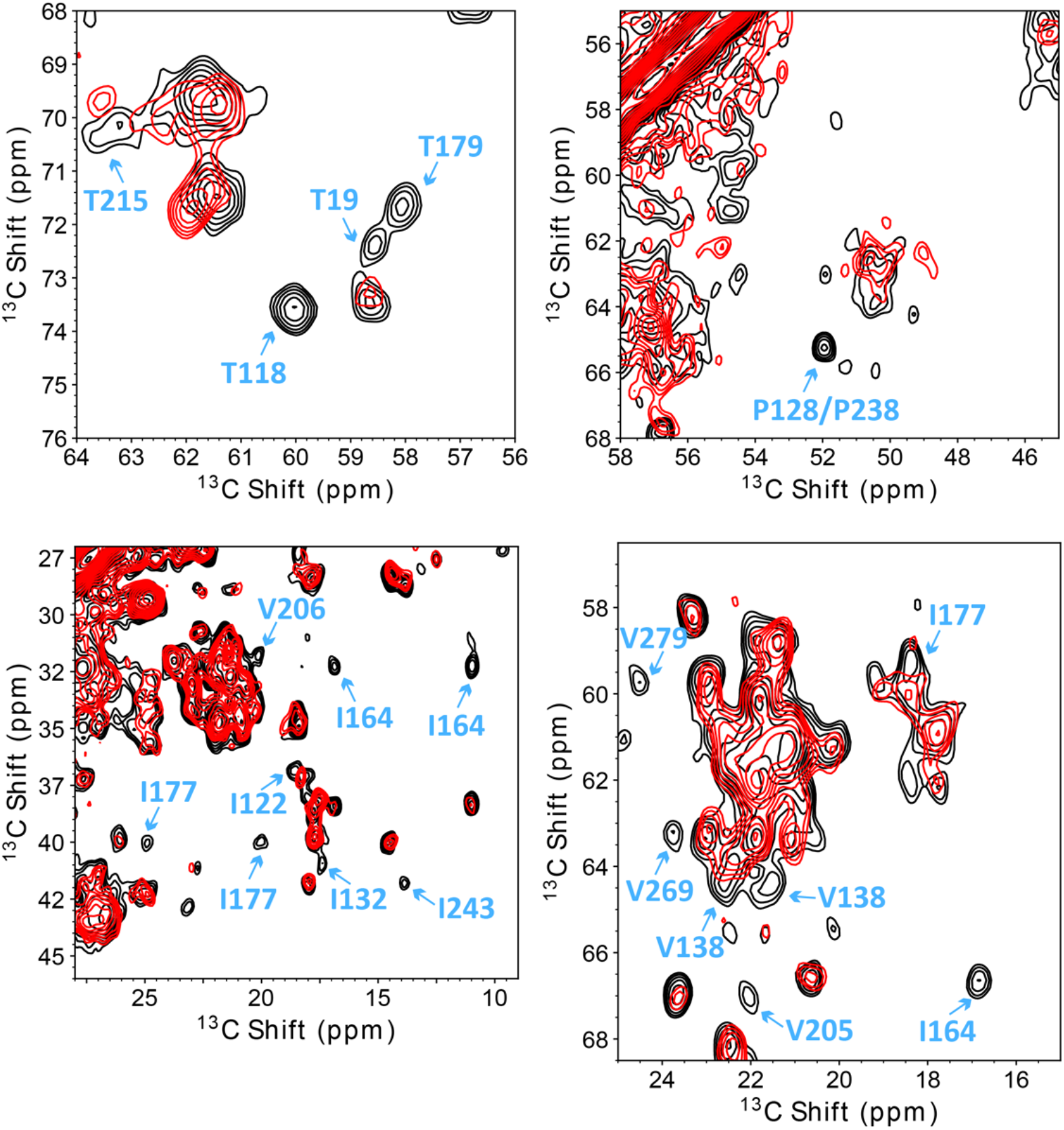
2D ^13^C homonuclear correlation spectra of fl TIA1:TC1 measured at 750MHz (black) and apo TIA1 (red) measured at 900MHz. The spectra were taken using the DARR(29) pulse sequence with a mixing time of 50 ms and a VT set temperature of 235K (actual sample temperature approximately 265K) and magic angle spinning frequency of 16.666 kHz. Lowest contours are at 3.75x RMS noise, and others are 1.25x higher. Blue arrows point to a few experimental peaks that change due to TC1 binding. These differences are mainly from side chain crosspeaks of threonine, isoleucine and valine residues in RRM2.

In figure 5 we display the 2D NCA spectra of fl apo TIA1 (red) and of fl TIA1 with 10nt TC1 ssDNA (black), and use labels to highlight various peaks that change due to DNA binding. Several examples are located in beta strands (V108, V137, V138) and loops (G173) of RRM2. The N-Cα peak of V108 appears in the spectrum of fl TIA1 with TC1, but not for apo fl TIA1. The N-Cα peak of V137 is shifted due to the presence of TC1. The N-Cα peak of V138 is dramatically reduced in intensity due to addition of TC1. Additional peaks in RRM3 are highlighted as well, which are located first alpha helix α_0_ (V205, V206), or in beta strands (I240, F253) and or in loops (G250, G276). The N-Cα peaks of V205 and V206 appear shifted and attenuated in the spectra of fl TIA1 with TC1 as compared with apo. The N-Cα peak of I240 does not appear in the presence of TC1. The N-Cβ peak of F253 appears only in the presence of TC1. The N-Cα peaks of G250 and G276 appear shifted in the presence of TC1. The perturbed peaks discussed above and displayed in figure 4 and 5 are summarized in Table 4 and highlighted in red color on the PDB structures of RRM2 and RRM3 in figure 6 and 7. Notably, these residues are all conserved in the sequence alignment (Fig. S3). Most of the side chain peaks that shift upon binding to TC1 are on the surface of the RRM2 domain. V108 and F253 are located in the highly conserved RNP motif.

**Table 4.**
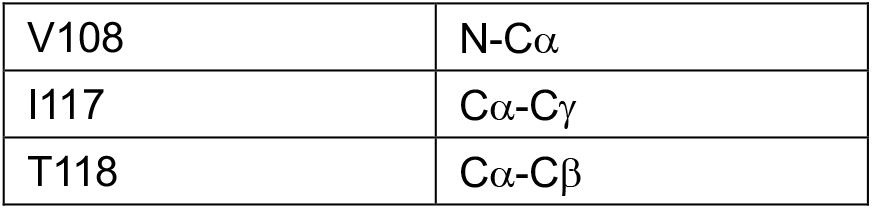
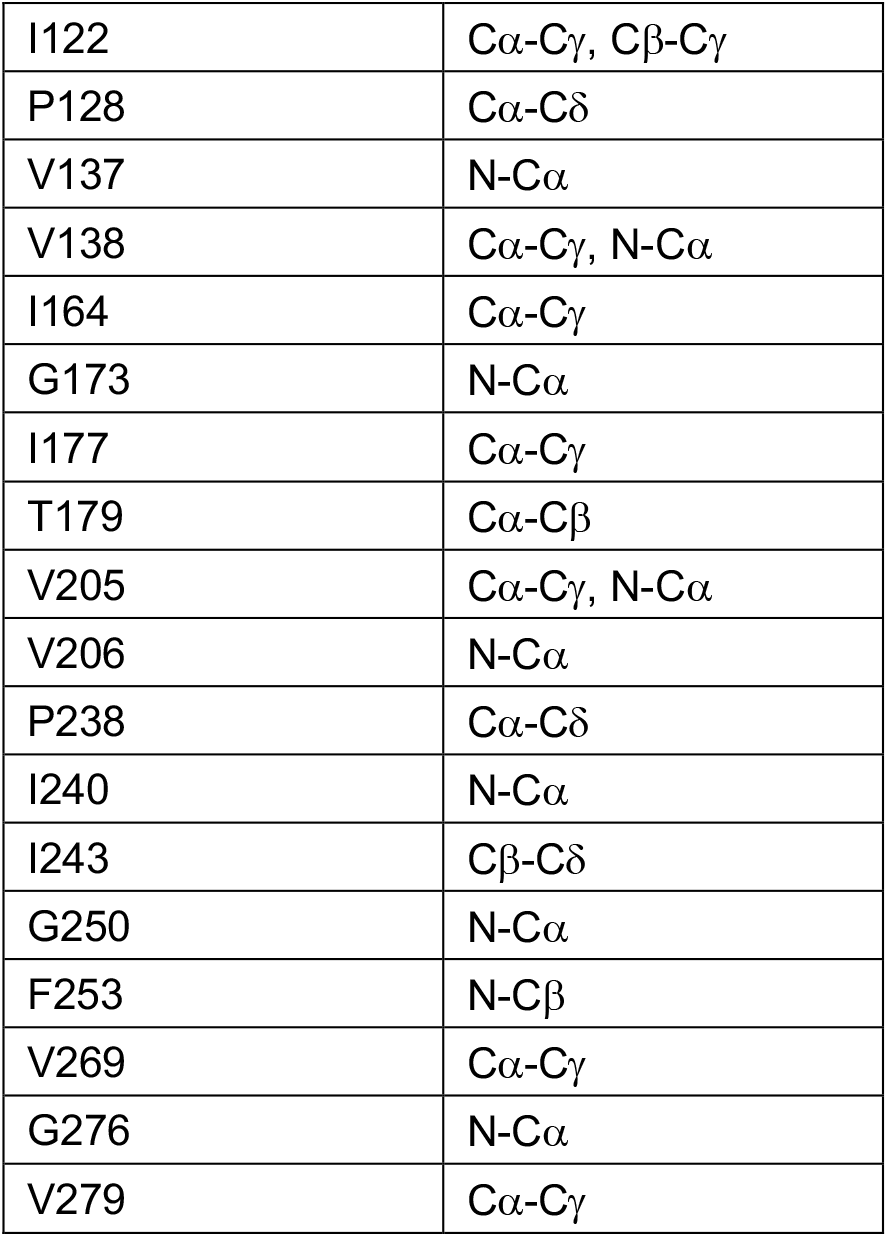
The residues with perturbed chemical shifts upon binding to TC1

|  |  |
| --- | --- |
| V108 | N-Cα |
| I117 | Cα-Cγ |
| T118 | Cα-Cβ |
| I122 | Cα-Cγ, Cβ-Cγ |
| P128 | Cα-Cδ |
| V137 | N-Cα |
| V138 | Cα-Cγ, N-Cα |
| I164 | Cα-Cγ |
| G173 | N-Cα |
| I177 | Cα-Cγ |
| T179 | Cα-Cβ |
| V205 | Cα-Cγ, N-Cα |
| V206 | N-Cα |
| P238 | Cα-Cδ |
| I240 | N-Cα |
| I243 | Cβ-Cδ |
| G250 | N-Cα |
| F253 | N-Cβ |
| V269 | Cα-Cγ |
| G276 | N-Cα |
| V279 | Cα-Cγ |

**Figure 5.**
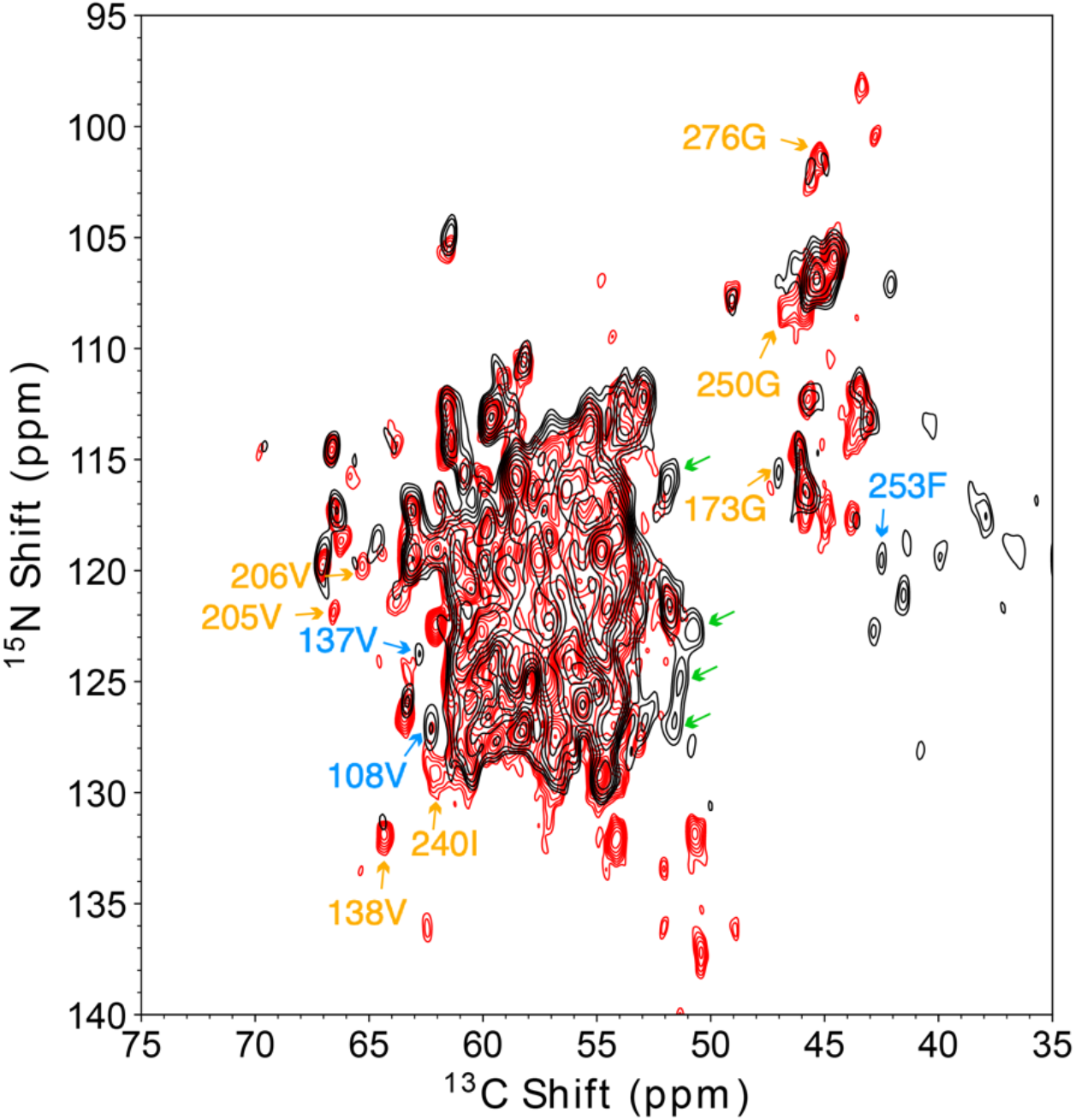
2D N-Cα heteronuclear correlation spectrum of fl TIA1:TC1 on 750MHz (black) and apo TIA1 (red) on 900MHz. The spectra were taken with double cross-polarization(38) pulse sequence at a set temperature of 235K (actual sample temperature approximately 280K) and the magic angle spinning frequency was 33.333 kHz. For NCA transfer, specific CP was optimized near the ν_1_(^13^C) =1.25 ν_r_ and ν_1_(^15^N) = 0.25 ν_r_ matching condition, with a 3.5 ms tangential ramp on the carbon channel. Lowest contours are at 4x RMS noise, and others are 1.25x higher. Peaks that change due to TC1 binding are pointed out with arrows. Peaks that appear only in the spectrum of fl TIA1:TC1 are highlighted with blue arrows while those seen only in the spectrum of apo TIA1 are highlighted with orange arrows. Peaks highlighted with green arrows are new peaks that appear in the spectrum of fl TIA1:TC1 but lack site specific assignments. (They are located relatively close to the chemical shifts of Pro Cδ.)

**Figure 6.**
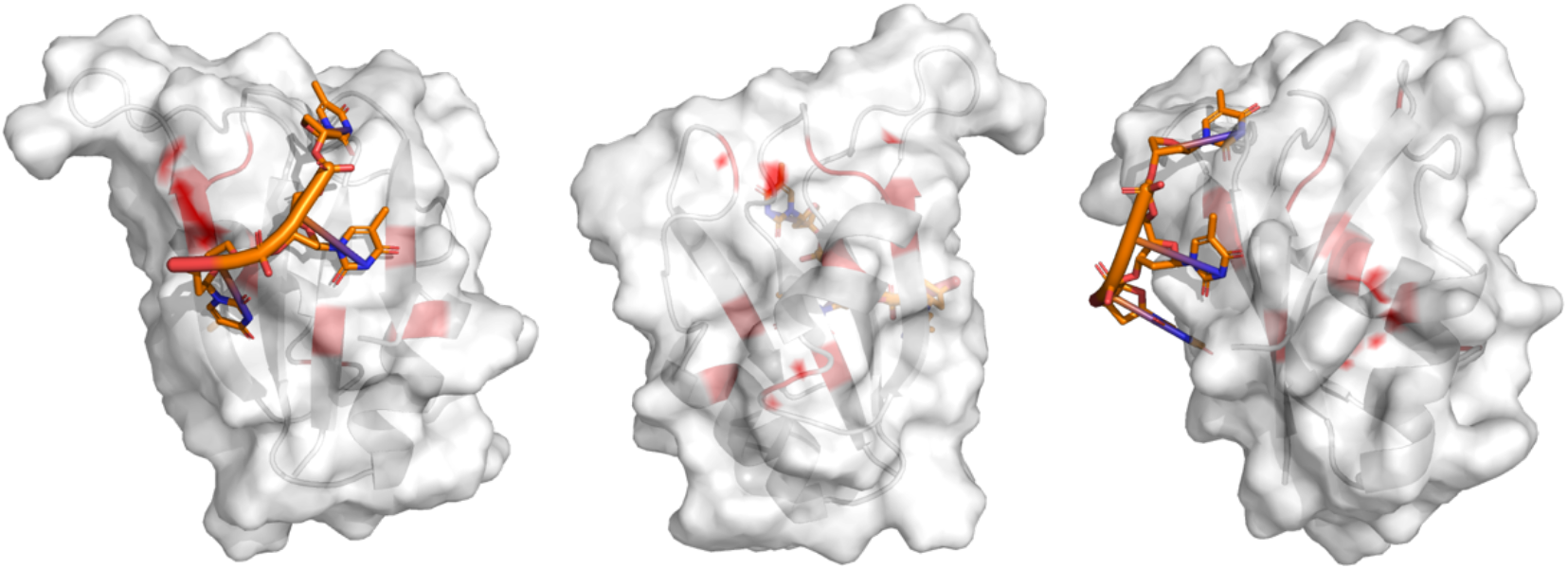
RRM2 homology model of mouse TIA1(25) aligned with crystal structure of RRM2 of human TIA1 in complex with ssDNA(24) (PDB: 5ITH, Organism: Homo sapiens, Method: X-ray diffraction, DNA chain: 5’-ACTCCTTTTT-3’, view from front, back, side direction). Residues in RRM2 exhibiting NMR chemical shift changes due to binding of TC1 ssDNA are highlighted on the structure in red. The red color appears on the surface are from side-chain changes and the red color appears on the secondary structure are from backbone changes.

**Figure 7.**
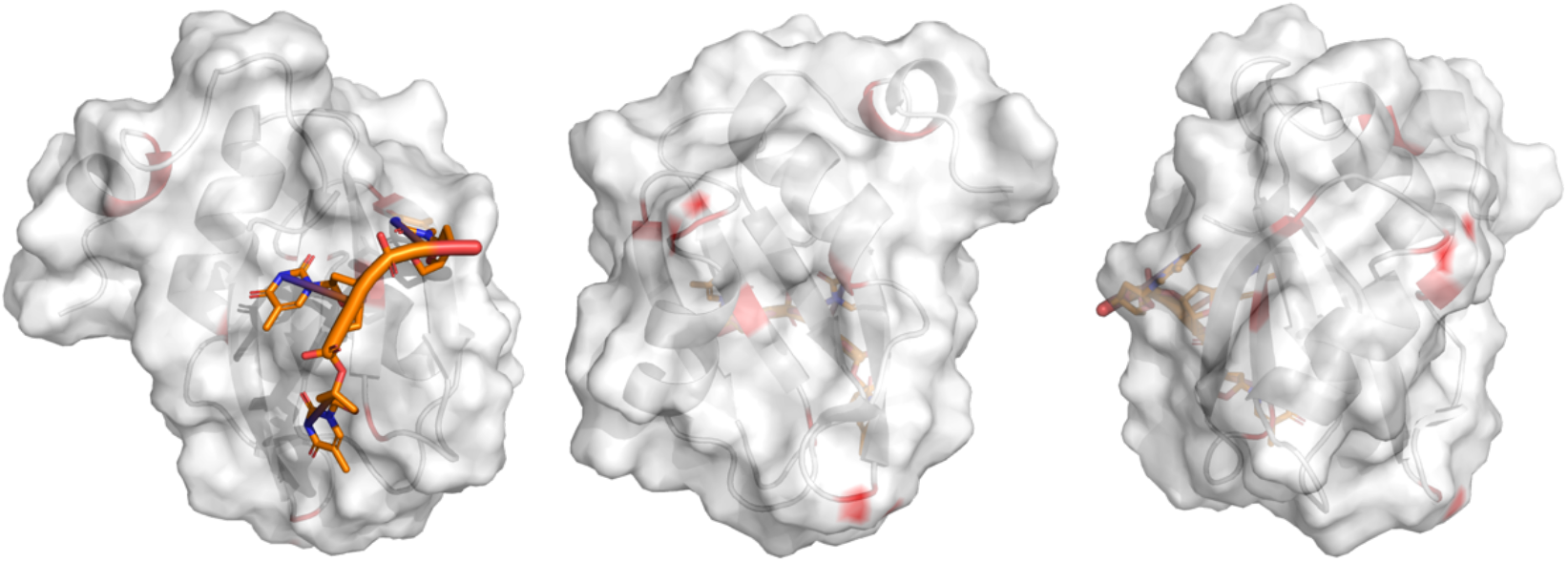
RRM3 homology model of mouse TIA1(25) aligned with crystal structure of RRM2 of human TIA1 in complex with ssDNA(24) (PDB: 5ITH, Organism: Homo sapiens, Method: X-ray diffraction, DNA chain: 5’-ACTCCTTTTT-3’, view from front, back, side direction). Residues in RRM3 exhibiting DNA induced NMR chemical shift changes are highlighted on the structure in red. The red color appears on the surface are from side-chain changes and the red color appears on the secondary structure are from backbone changes.

Our NMR shift perturbation studies clarify the nucleic acid binding behavior of oligomeric fl TIA1, and are in good agreement with prior studies of nucleic acid interactions. The residues with perturbed chemical shifts in RRM2 (highlighted in Fig 6) were relatively close to the nucleic acid binding surfaces identified in previous structural studies (PDB: 5ITH, Method: X-ray diffraction)(24), while the few perturbed residues in RRM3 were not close to previously identified binding regions (Fig 7).

Assuming that chemical shift perturbations are indicators of the nucleic acid binding surface, we conclude that TC1 binds to RRM2 and not to RRM3 in the oligomeric form under our experimental conditions. This is expected base on the fact that it is T-rich and thus designed to bind to RRM2. The few perturbations in RRM3 presumably result from the 3’-end of TC1 touching RRM3 tentatively or small conformational changes and interactions between RRM2 and RRM3 coupled to TC1 binding. This conclusion also agrees well with the results of isothermal titration calorimetry experiments that indicated that TC1 binds tightly to TIA1 with a 1:1 stoichiometry (and only weakly at a second site).

### Comparison of Zn^2+^ bound and apo forms of TIA1 by SSNMR

In figure 8 we display the 2D ^13^C-^13^C correlation spectra of fl TIA1 (red) and of fl TIA1 with Zn^2+^ (black). The side chain crosspeaks (Cα-Cγ, Cb-Cγ, Cγ_1_-Cγ_2_) of T119, I122, I164, I177 in RRM2 and V205, K280 in RRM3 appear in the presence of Zn^2+^, but not in the absence, and Cα-Cβ crosspeaks of T118 in RRM2 and T215, T222 in RRM3 increased in intensity due to addition of Zn^2+^. On the other hand, C-Cβ peak of T179 and Cα-Cβ peak of A188 in the linker region between RRM2 and RRM3 that appear in the TIA1 spectrum are missing in the presence of Zn^2+^. Additionally, several crosspeaks for residues in RRM3 do not appear in the presence of Zn^2+^, including Cα-Cβ peaks of Y202 and T226, Cα-Cγ peak of T233, and Cβ-Cγ peak of V221.

**Figure 8.**
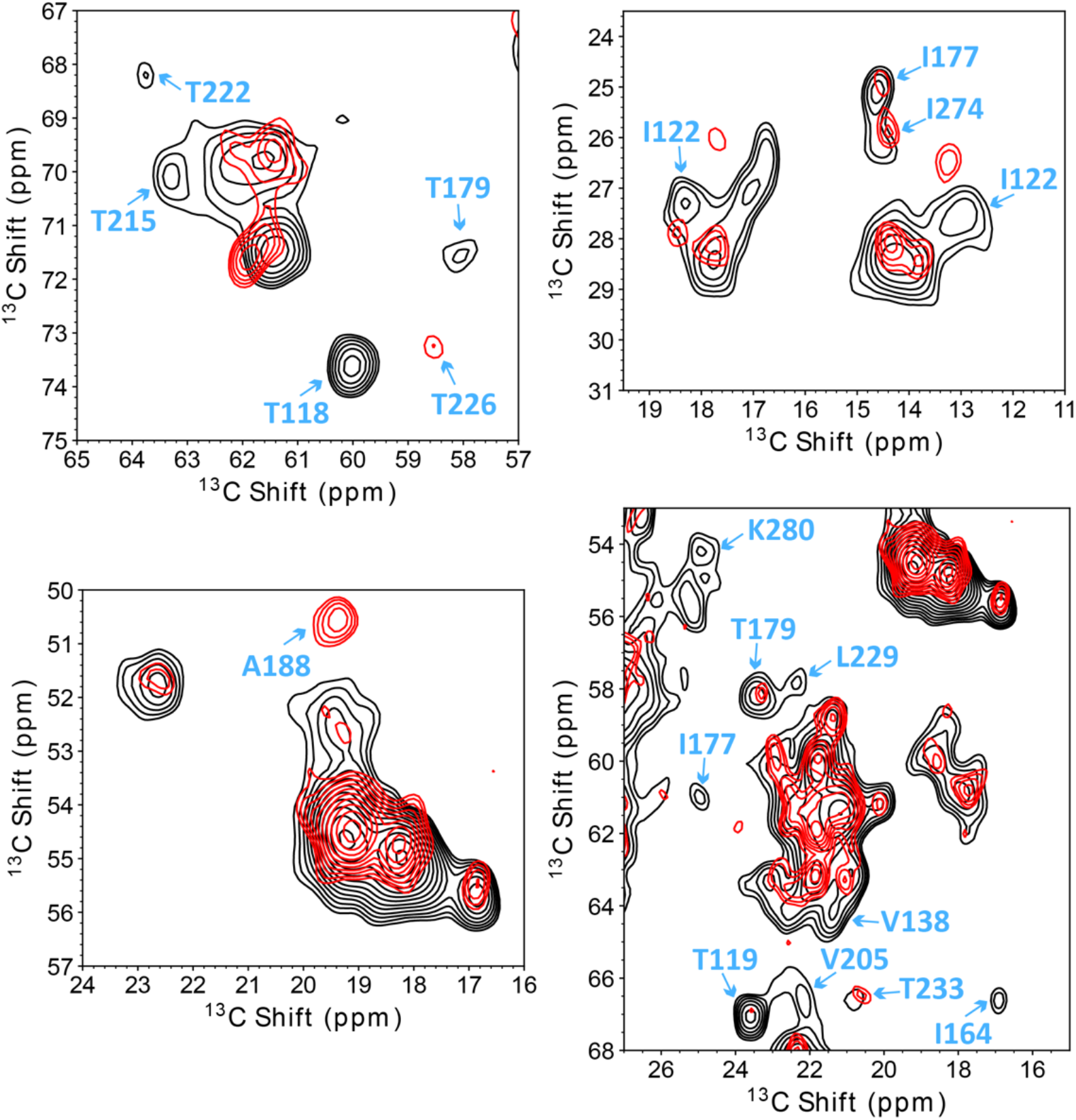
2D ^13^C homonuclear correlation spectrum of fl TIA1: Zn^2+^ on 750MHz (black) and apo TIA1 (red) on 900MHz. The spectra were taken using the DARR pulse sequence with a VT set temperature of 235K (actual sample temperature was around 265K) and the magic angle spinning frequency was 16.666 kHz. The mixing time was 50 ms. Lowest contours are at 3.75x RMS noise, and others are 1.25x higher. Arrows point to a few peaks that change due to Zn^2+^ binding.

In figure 9 we display the 2D NCA spectra of fl TIA1 (red) and of fl TIA1 with Zn^2+^ (black). Labels highlight several peaks that change due to Zn^2+^ binding. The N-Cβ peak of T119 and S146 in RRM2 appears in the spectrum of fl TIA1 with Zn^2+^, but not for apo fl TIA1. The N-Cα peak of G168 in RRM2 and V205, V206 in RRM3 are shifted due to the presence of Zn^2+^. Additionally, the N-Cα peak of G276 in RRM3 is missing when TIA1 binding with Zn^2+^.

**Figure 9.**
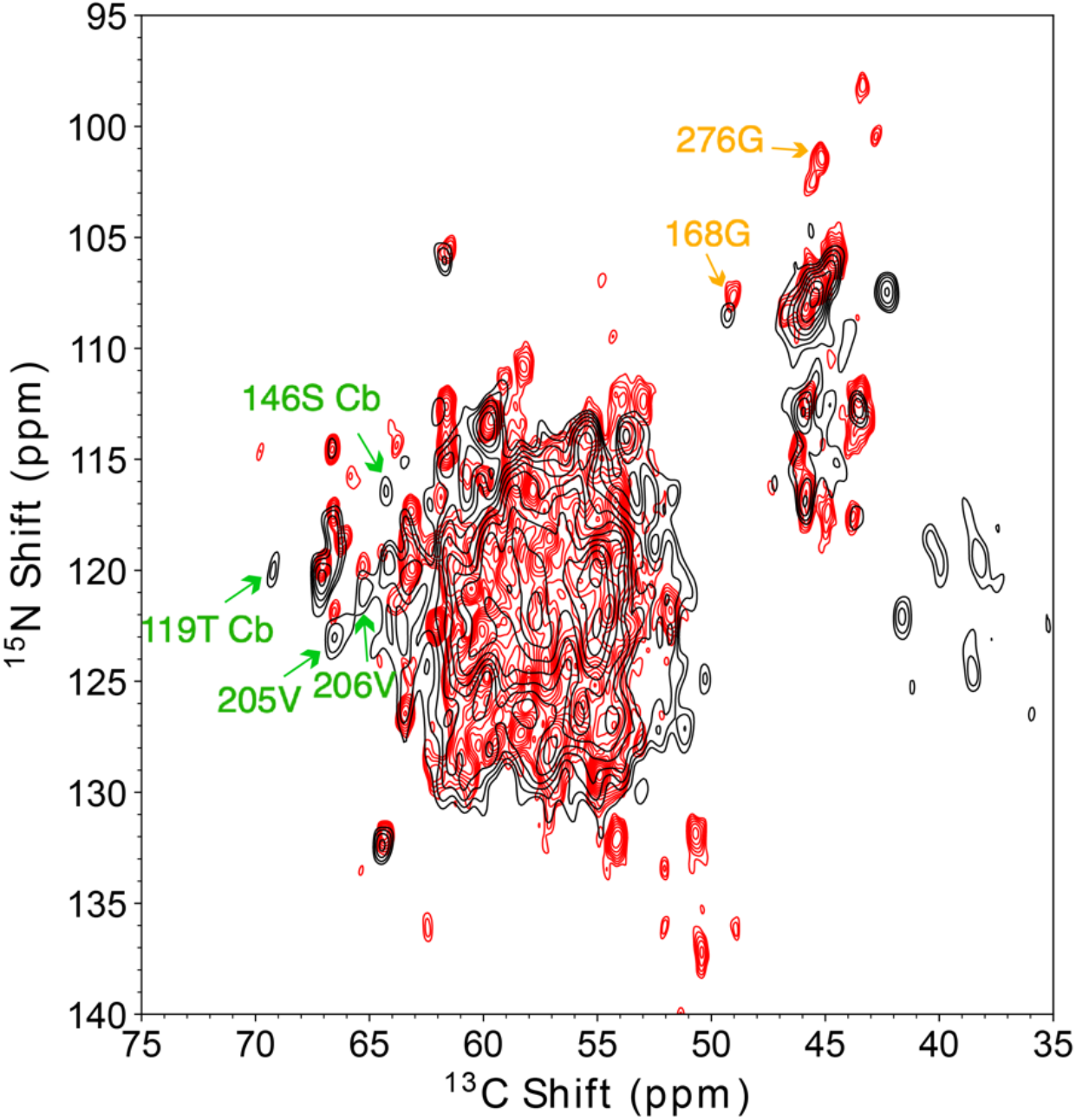
2D N-Cα heteronuclear correlation spectrum of fl TIA1:Zn^2+^ on 750MHz (black) and apo TIA1 (red) on 900MHz. The spectra were taken with double cross-polarization pulse sequence at a set temperature of 235K (sample temperature was around 280K) and the magic angle spinning frequency was 33.333 kHz. For NCA transfer, specific CP was optimized near the ν_1_(^13^C) =1.25 ν_r_ and ν_1_(^15^N) = 0.25 ν_r_ matching condition, with a 3.5 ms tangential ramp on the carbon channel. Lowest contours are at 4x RMS noise, and others are 1.25x higher. Peaks that change due to Zn^2+^ binding are pointed out with arrows. Peaks appears only in the spectrum of fl TIA1 with Zn^2+^ are pointed out with green arrows while only in the spectrum of apo TIA1 are pointed out with orange arrows.

The Zn^2+^ binding sites of RRM domains were predicted using the Metal Ion-Binding Site Prediction and Docking Server(31), which identifies local structure motifs based on a structural alignment algorithm that combines both structure and sequence information (Fig. 10). There are two partial Zn^2+^ binding sites in RRM2. One site mainly consists of H105, H107, with D134 probably involved as well. The other site is composed of E116 and K145. There are three partial Zn^2+^ binding sites in RRM3, which are (1) H259, E260; (2) E260, H264; and (3) C218, C281. Residues that experience chemical shift perturbations (in either the DARR or the NCA spectra) upon binding to Zn^2+^ are shown as sticks and highlighted in red (Fig. 10). For the perturbed residues in RRM2, T118 and S146 are close to the predicted Zn^2+^ binding half site composed of E116 and K145. Apart from them, other residues in RRM2 that show chemical shift perturbation with the addition of Zn^2+^ are all far from the predicted Zn^2+^ binding sites. For residues in RRM3 that exhibit chemical shift perturbations upon binding to Zn^2+^, V221, T222, T233, and K280 are close to the predicted half site composed of C218 and C281; Y202 and V206 and are close to the predicted half site composed of H259 and E260; Y202 is close to the predicted half site composed of E260 and H264 as well. Therefore, the data suggest that RRM3 probably interacts directly with Zn^2+^ as predicted and all of the three predicted Zn^2+^ binding half site interact with and ligate Zn^2+^, while only one of the predicted Zn^2+^ binding half site in RRM2 interacts with Zn^2+^. Possibly the unexpected pattern in chemical shift perturbation for RRM2 are due to small conformation changes upon Zn^2+^ binding without any indication of secondary structure changes.

**Figure 10.**
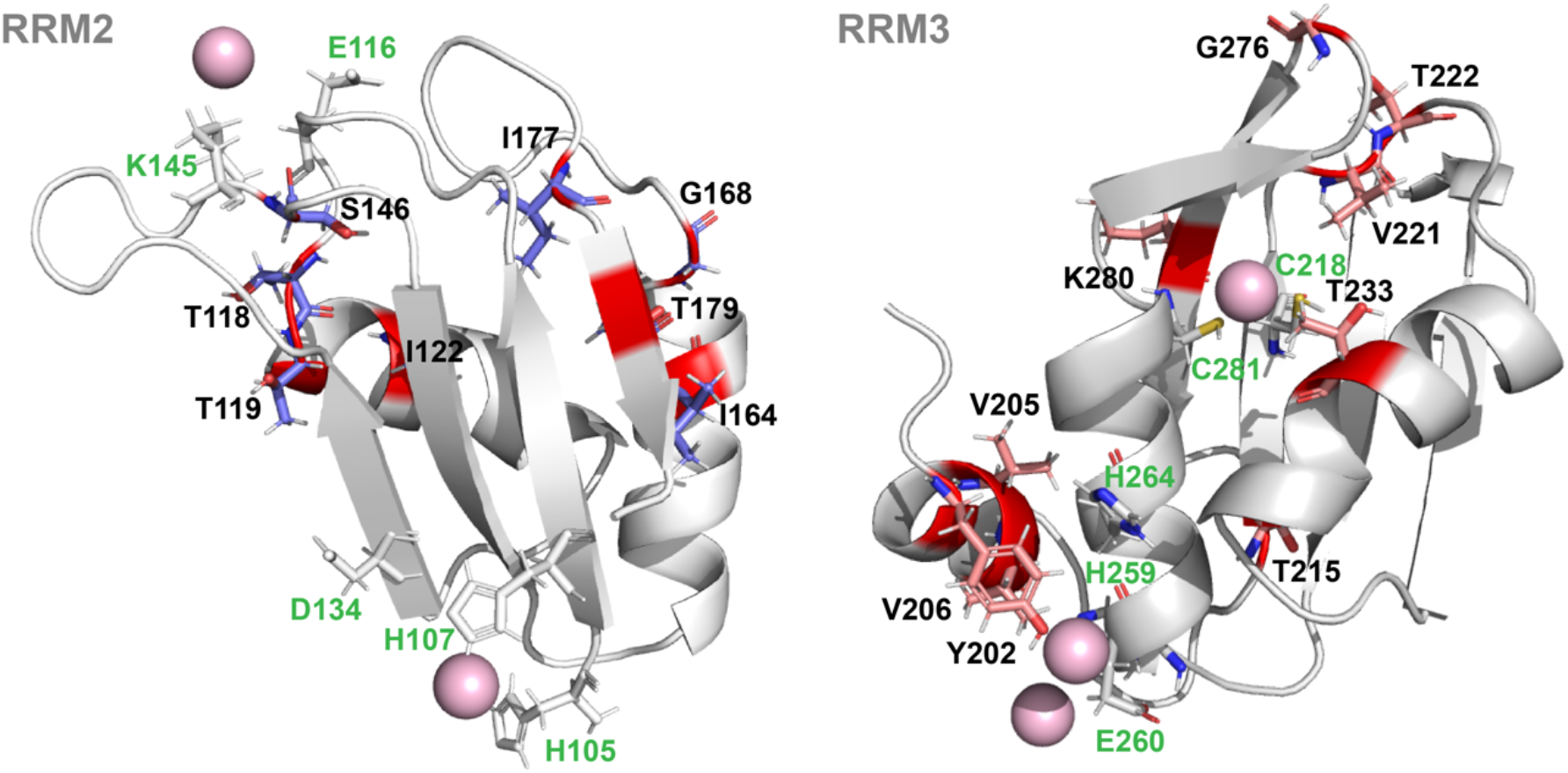
Shifted residues in DARR and NCA spectra due to Zn^2+^ binding highlighted on the RRM2 and RRM3 structures. The pink spheres represent expected Zn^2+^ locations (based on predictions using Metal Ion-Binding Site Prediction and Docking Server). The predicted Zn^2+^ binding sites are all half sites. There are two half sites in RRM2. One half site mainly consists of H105, H107, with D134 probably involved as well. The other half site is composed of E116 and K145. There are three half sites in RRM3, which are (1) H259, E260; (2) E260, H264; and (3) C218, C281. These residues related to predicted half sites are shown as white sticks and marked with green texts. By contrast, the residues that are identified with sequence numbers (black) show chemical shift perturbations upon binding of Zn^2+^ and are shown as sticks (purple in RRM2 and salmon in RRM3) and highlighted in red color. For the perturbed residues in RRM2, T118 and S146 are close to the predicted Zn^2+^ binding half site composed of E116 and K145. For the perturbed residues in RRM3, V221, T222, T233, and K280 are close to the predicted half site composed of C218 and C281; Y202 and V206 and are close to the predicted half site composed of H259 and E260; Y202 is close to the predicted half site composed of E260 and H264 as well.

### Simultaneous binding of TC1 and Zn2_+_

In figure 11 we display the 2D ^13^C homonuclear correlation spectrum of fl TIA1 with TC1 (black) and TIA1 with both TC1 and Zn^2+^ (red), and use labels to highlight various peaks that change due to binding. The side chain peaks of I39, I132, L229, I240, I243, I266 appear in the doubly bound species, whereas they are not observed in the spectrum of TIA1 with only TC1 bound. The Cα-Cβ, Cβ-Cγ, of T19 and Cβ-Cγ peak of I122 are missing in presence of both TC1 and Zn^2+^. An unassigned Ala Cα-Cβ peak also appears in the spectrum of TIA1 with only TC1. Based on the chemical shifts, it is likely to be in a “random coil” conformation, and is possibly from RRM1.

**Figure 11.**
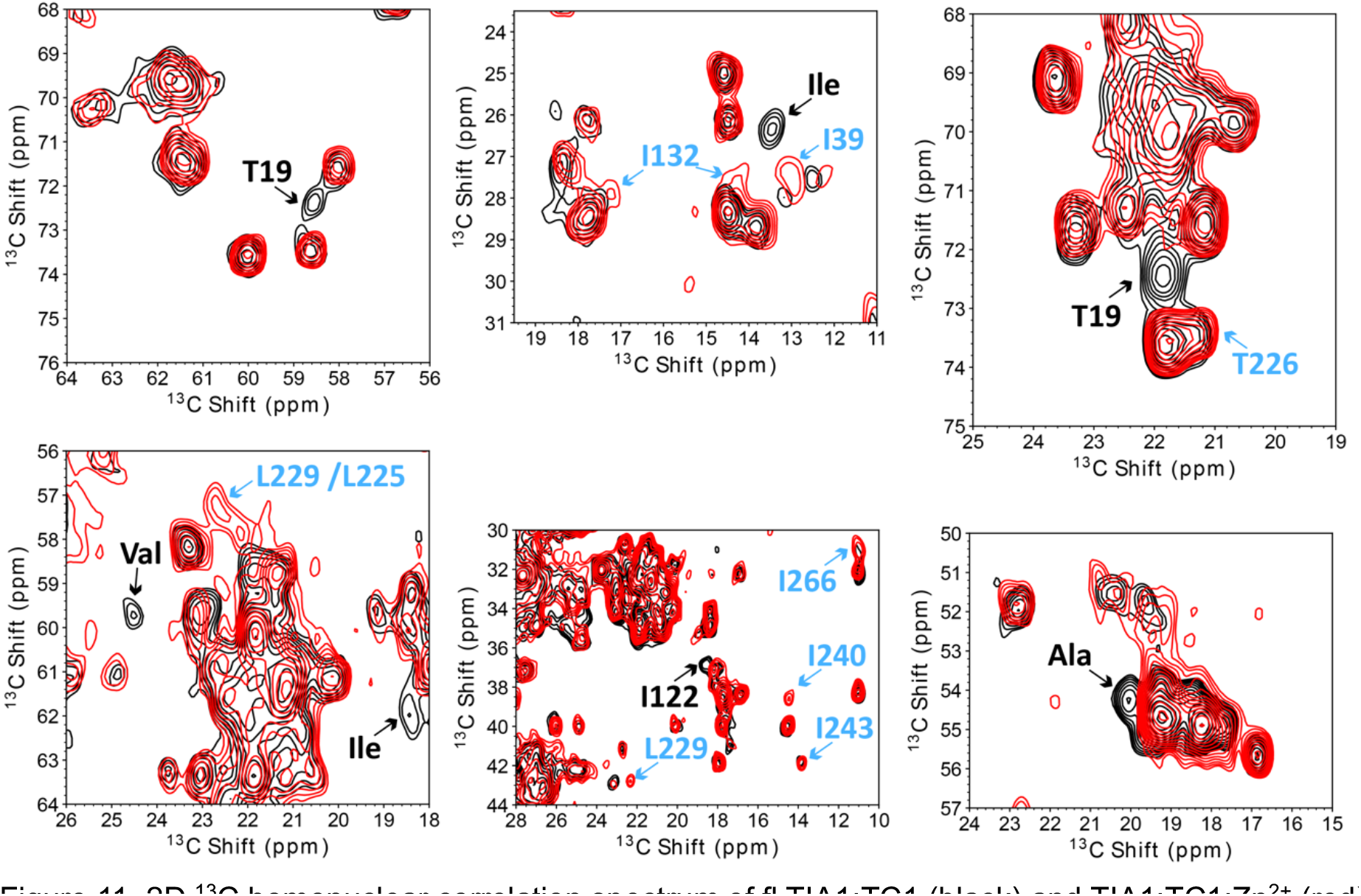
2D ^13^C homonuclear correlation spectrum of fl TIA1:TC1 (black) and TIA1:TC1:Zn^2+^ (red) measured at 750MHz. The spectra were taken with the DARR pulse sequence (mixing time 50 ms) at a set temperature of 235K (actual sample temperature was approximately 265K) and the magic angle spinning frequency was 16.666 kHz. Lowest contours are at 3.75x RMS noise, and others are 1.25x higher. Arrows point to a few peaks that change due to Zn^2+^ and DNA binding. These differences are mainly from side chain crosspeaks of isoleucines.

In figure 12 we display the NCA spectra of TIA1 with different ligands. Peaks that appear in the presence of TC1 are highlighted with blue arrows, while peaks that appear in presence of Zn^2+^ are highlighted with purple arrows. The peaks that appear only in presence of both TC1 and Zn^2+^ are highlighted with orange arrows. These residues could be assumed to be located in the binding site. For example, the N-Cβ peak of S146 appears on both spectra when Zn^2+^ is added but does not appear on TIA1 with TC1 nor on apo TIA1, and therefore S146 is assumed to be perturbed specifically due to Zn^2+^ binding. The N-Cα peak of V108 appears in both spectra of TIA1:TC1 but not in spectra of TIA1:Zn^2+^ nor apo TIA1, so we conclude that V108 is specifically perturbed due to DNA binding. However, the N-Cα peak of V138 is missing in TIA1:TC1 but appears is the other three conditions (TIA1, TIA1 with Zn^2+^, TIA1 with both TC1 and Zn^2+^), indicating that Zn^2+^ binding sites could overlap with DNA binding sites or somehow allosterically be connected. From the back-side views of RRM2 and RRM3 (Fig. 13 and 14), there are also some residues affected by TC1 binding which could result from indirect effects such as allosteric changes.

**Figure 12.**
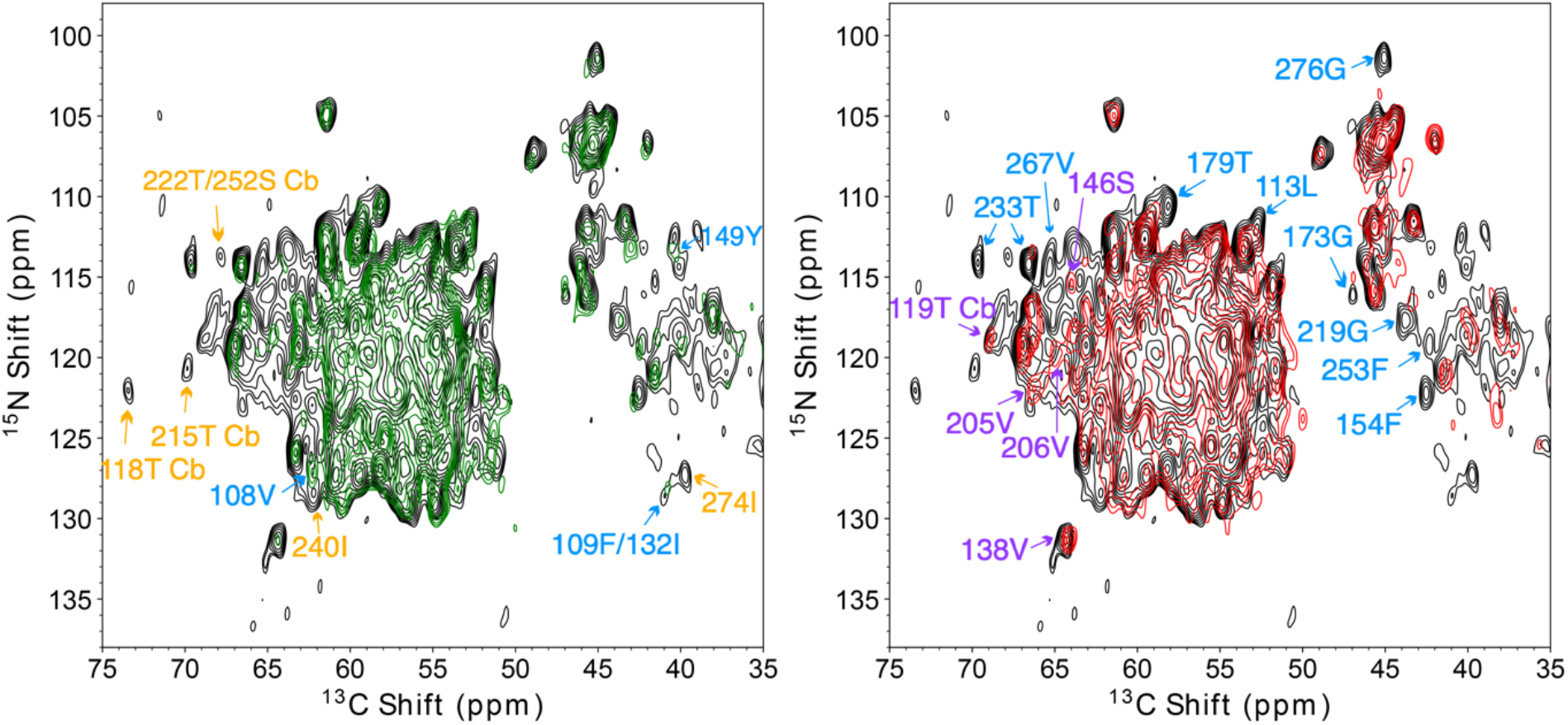
2D N-C heteronuclear correlation spectrum of fl TIA1:TC1 (green), TIA1:Zn^2+^ (red) and TIA1:TC1:Zn^2+^ (black) on 750MHz. The spectra were taken with double cross-polarization pulse sequence at a set temperature of 235K (sample temperature was around 280K) and the magic angle spinning frequency was 33.333 kHz. For NCA transfer, specific CP was optimized near the ν_1_(^13^C) =1.25 ν_r_ and ν_1_(^15^N) = 0.25 ν_r_ matching condition, with a 3.5 ms tangential ramp on the carbon channel. Lowest contours are at 4x RMS noise, and others are 1.25x higher. The blue residue labels indicate peaks observed in the presence of TC1 (green and black) but not in the absence (red). The purple residue labels residues indicate residues whose peaks are observed shown in the presence of Zn^2+^ (red and black) but not in its absence (green). The orange residue labels indicate residues for which the peaks are only observed when both Zn^2+^ and TC1 are bound (black).

**Figure 13.**
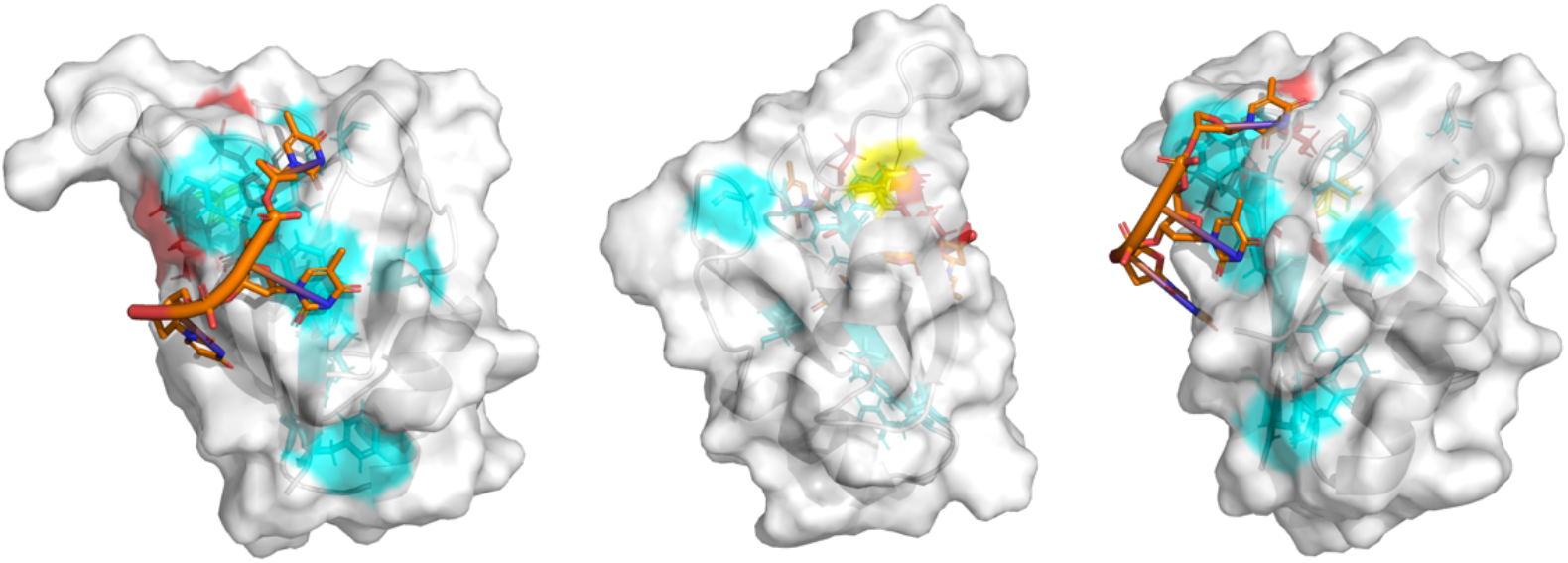
Structure of RRM2 with regions that are affected by TC1 binding (blue), Zn^2+^ binding (red) and both TC1 and Zn^2+^ binding (yellow) based on chemical shift perturbations.

**Figure 14.**
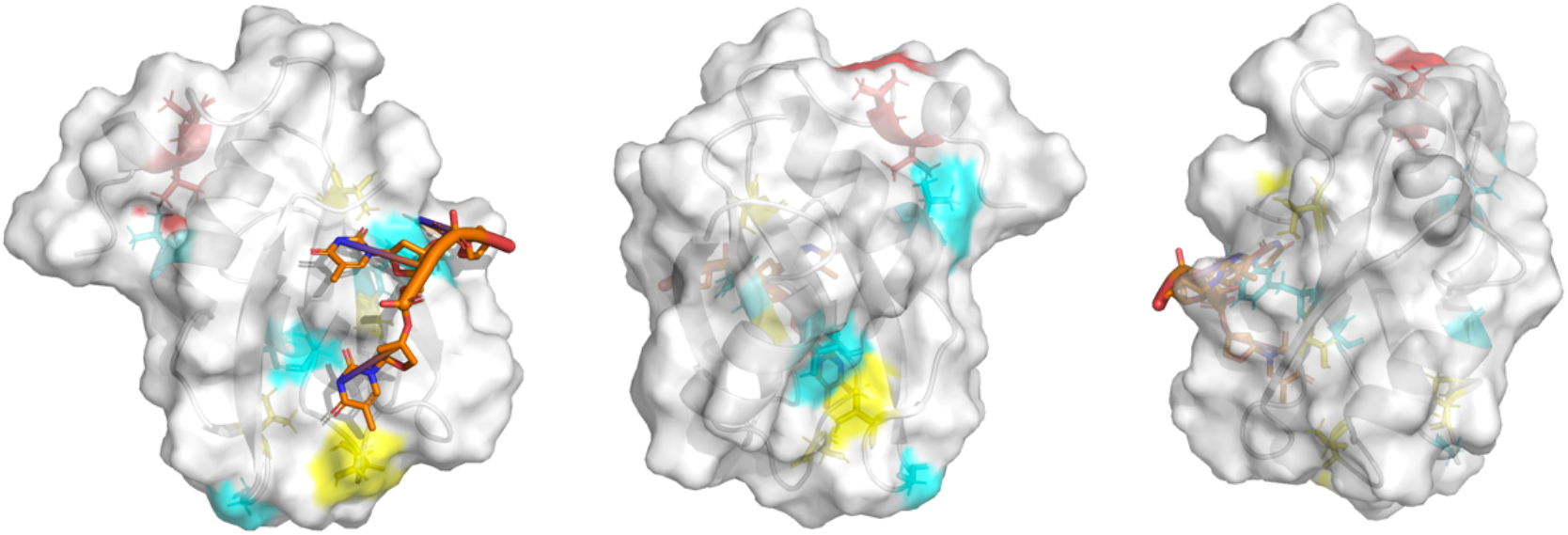
Structure of RRM3 with regions that affected by TC1 binding (blue), Zn^2+^ binding (red) and both TC1 and Zn^2+^ binding (yellow) based on chemical shifts perturbations.

Apart from the residues in RRM2 and RRM3 domains, the new peaks that appear in the DARR spectrum of TIA1 with TC1 (Fig. 11) in the Threonine Cα-Cβ and Cβ-Cγ region could be assigned to T19 in RRM1, since the predicted shifts for this residue are Cα at 59.24 ppm, Cβ at 72.19 ppm and Cγ_2_ at 21.43 ppm. Most other residues in RRM1 appear to remain undetectable, presumably due to conformational dynamics. Based on the RRM1 and RRM2 docking result using HADDOCK server(32) with default parameters and homology models of RRM1 built from PDB: 2CQI and RRM2 built from PDB:2MJN (25) (Fig. 15), T19 in RRM1 is located in the RRM1 and RRM2 interface in the most confidently predicted cluster of the protein domain docking model. Possibly, the DNA and its interactions with the protein has the effect of aligning the RRM domains, giving RRM1 a preferential orientation with respect to RRM2, resulting in T19 being detected. RRM1 potentially assists DNA binding (but not Zn^2+^) by providing a net positively charged environment surround the DNA.

**Figure 15.**
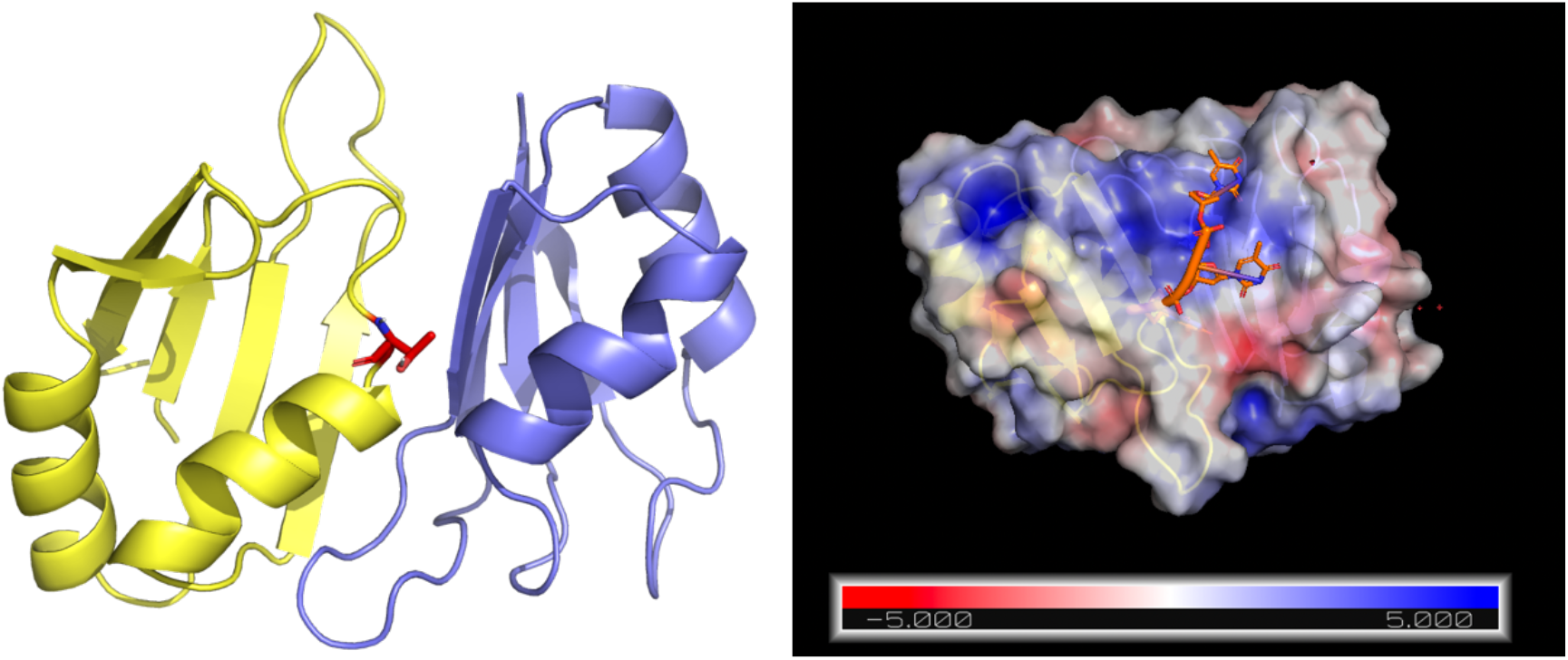
The top cluster of RRM1 (yellow) and RRM2 (purple) docking models using HADDOCK server. 19T in RRM1 is shown as sticks and highlighted in red color. The electrostatic surface of RRM12 is shown on the right side. The Cα -Cβ, Cβ-Cγ of T19 in RRM1 are newly detected when TIA1 bound to TC1, indicating a potential surface where interdomain interactions presents between RRM1 and RRM2 when TC1 bound to RRM2. The positively charged environment make the DNA bound to RRM2 more stable.

### TIA1 bound to single strand DNA with multiple binding sites

Although DNA (TC1-A_8_)_2_-TC1 and TC1 have very different effects on inducing the oligomerization of TIA1 in a turbidity assay, the NMR spectra of TIA1: (TC1-A_8_)_2_-TC1 closely resemble those for TIA1:TC1, apart from a few differences. One of the differences is that the crosspeaks (Cα-Cβ, Cβ-Cγ, Cγ-Cδ) of I164 is missing on spectrum of TIA1: (TC1-A_8_)_2_-TC1 in comparison of with TIA1:TC1 (Fig. 16), as is the N-Ca peak of I164 in the NCA spectrum (Fig. 17). I164 may be located on the interface of TIA1 monomers. (TC1-A_8_)_2_-TC1 possibly brings the monomers together upon binding and affect the rigidity of the residues at that interface. In conclusion, there is no apparent difference in the protein-DNA binding interface comparing the two DNA constructs, suggesting that TIA1 has a similar conformation when bound to DNA with one vs multiple binding sites, and the multivalency of the larger DNA does not dramatically rearrange TIA1. Their different behaviors in promoting oligomerization can be attributed to that DNA with multiple binding site tends to link the monomers together more spontaneously even without agitation.

**Figure 16.**
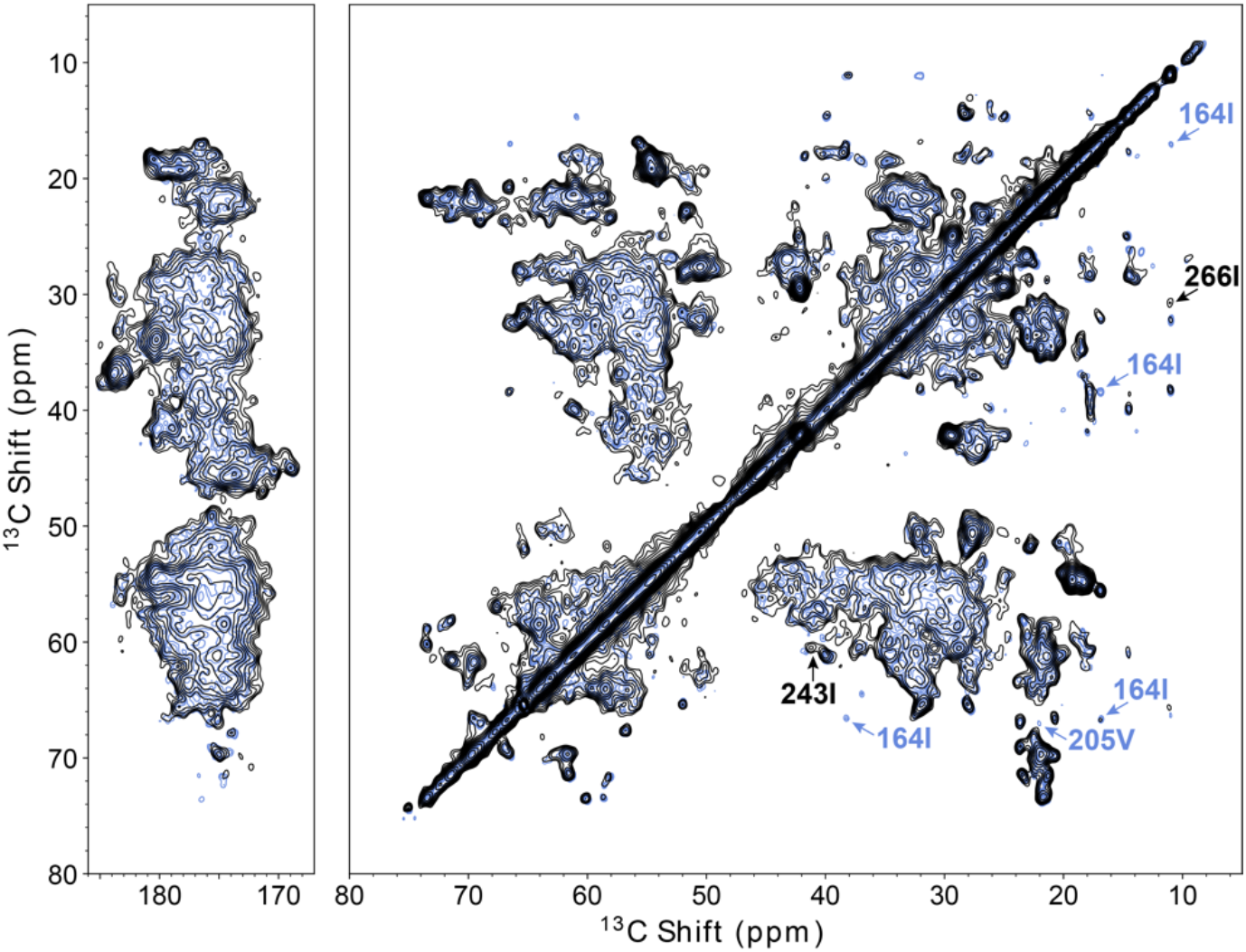
2D ^13^C homonuclear correlation spectrum of fl TIA1 with TC1 (blue) and with (TC1-A_8_)_2_-TC1 (black) on 750MHz. The spectra were taken with DARR pulse sequence at a set temperature of 235K (sample temperature was around 265K) and the magic angle spinning frequency was 16.666 kHz. The mixing time was 50 ms. Lowest contours are at 3.75x RMS noise, and others are 1.25x higher. The DNA sequence of TC1 was 5’-TTTTTACTCC-3’. The DNA sequence of (TC1-A_8_)_2_-TC1 was 5’-TTTTTACTCC AAAAAAAA TTTTTACTCC AAAAAAAA TTTTTACTCC-3’. It is a duplicate of TC1 with A linkers between each binding site. The arrows in the spectrum point to a few experimental peaks that change due to different DNA binding. Apart from peaks of I164, I243, I266 and V205 that pointed out with arrows, there are no other differences.

**Figure 17.**
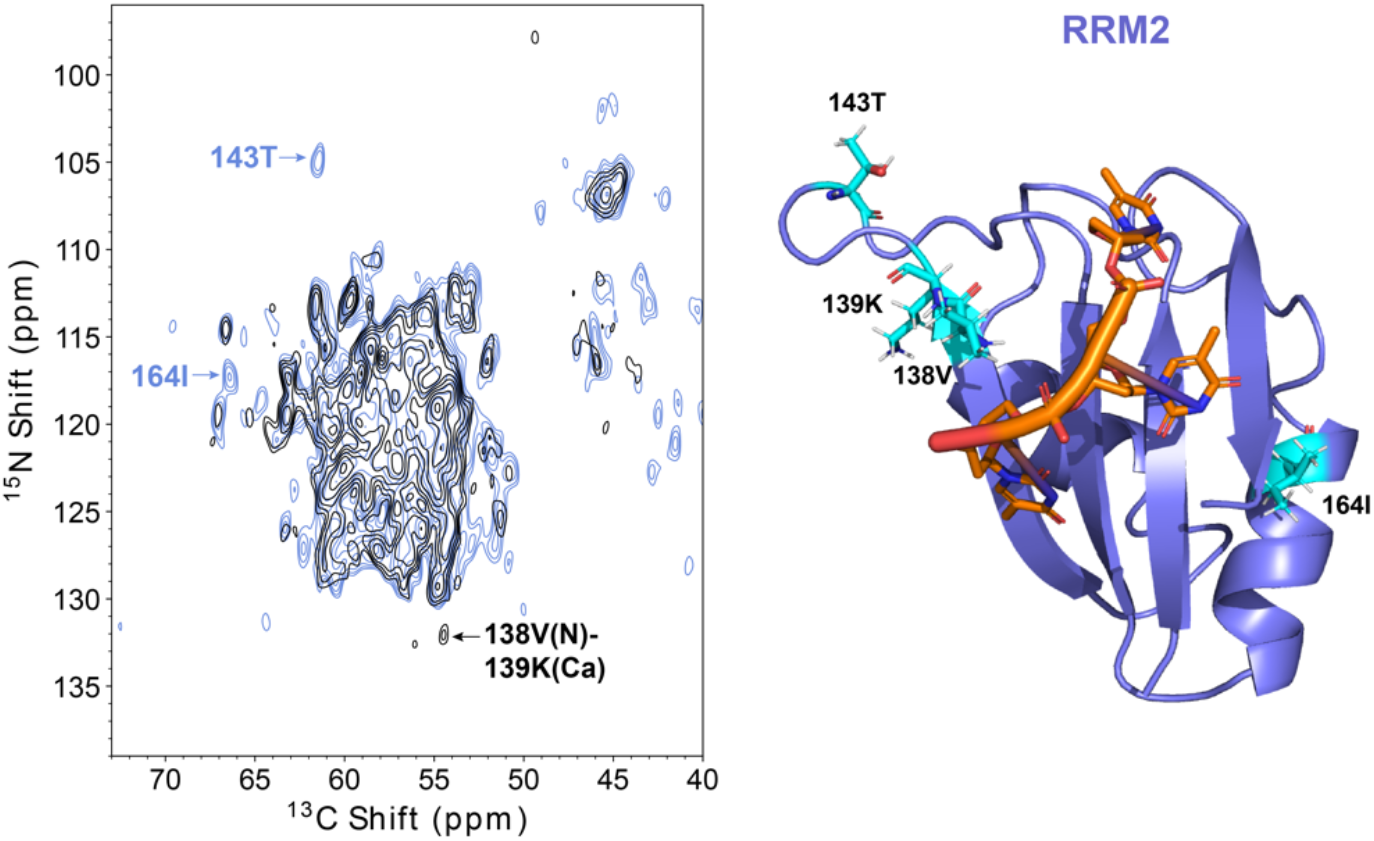
2D N-C heteronuclear correlation spectrum of fl TIA1 with (TC1-A_8_)_2_-TC1 on 750MHz (black) and TIA1 with TC1 on 750MHz (blue). The spectra were taken with double cross-polarization pulse sequence at a set temperature of 235K (sample temperature was around 280K) and the magic angle spinning frequency was 33.333 kHz. For NCA transfer, specific CP was optimized near the ν_1_(^13^C) =1.25 ν_r_ and ν_1_(^15^N) = 0.25 ν_r_ matching condition, with a 3.5 ms tangential ramp on the carbon channel. Lowest contours are at 4x RMS noise, and others are 1.25x higher. Peaks that change due to different DNA binding are pointed out with arrows. The homology model of RRM2 (purple) was shown in complex with DNA (orange) (DNA sequence: 5’-ACTCCTTTTT-3’). Residues experience chemical shifts changes when bound to (TC1-A_8_)_2_-TC1 were shown as sticks in cyan.

### Supramolecular models of oligomeric TIA1

Based on the EM images and NMR results, we propose models for oligomeric TIA1 with various ligands (Fig. 18). Panel A shows the domain structure of fl TIA1 monomer. Panel B shows a model of oligomeric apo TIA1. The oligomerization of apo TIA1 is likely to be promoted by interactions between PRDs from different TIA1 monomers. Agitation accelerates this process without changing the final morphology of apo TIA1 oligomers. Panel C shows a model of DNA (TC1) bound TIA1 oligomers. TC1 presumably acts as a scaffold, linking the monomers by binding to RRMs from different monomers, and forming oligomers appearing as long strings (Fig. 3B and 18C). For single stranded DNA, the theoretical effective length (rise) of each nucleotide is 0.7 nm, so for a ten-mer such as TC1 the total length is up to 7 nm. Assuming a 2∼3 nm diameter for each RRM, a 10nt DNA might be able to link 2 or 3 RRM domains. Given its sequence and binding properties, TC1 might link RRM2 and RRM3 from the same monomer, or it might link RRMs from different monomers. Without agitation, TC1 does not induce or accelerate oligomerization, therefore under quiescent conditions, it is reasonable to assume that it might only link RRM2 and RRM3 from the same monomer. Following agitation, oligomers form, with distinct EM images. In the TIA1:TC1 complex the molecules seem to be linked into a larger mesh, which supports the suggestion that TC1 binds preferentially links to RRMs from multiple monomers to form a “daisy chained” larger structure.

**Figure 18.**
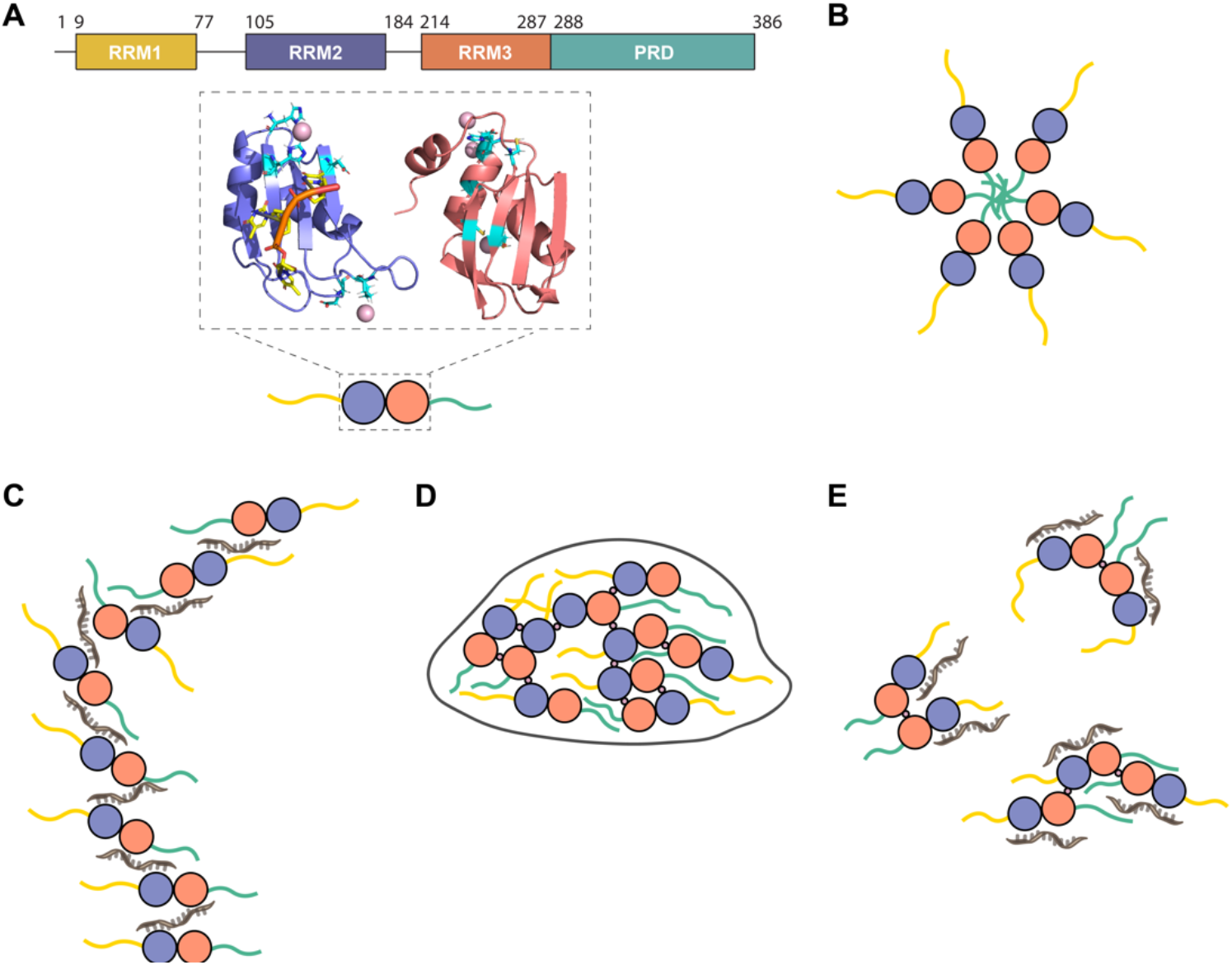
(A) Domain structures of TIA1-RRM2 (purple) and RRM3 (salmon) with predicted Zn^2+^ binding sites (light green). RRM1 (yellow) and PRD (dark green) domains are disordered, therefore shown as lines. Zn^2+^ are shown as pink spheres and the highlighted residues are predicted to be involved in Zn^2+^ binding are in light green. (B-E) Oligomeric models of (B) oligomeric apo fl TIA1, (C) TIA1 with TC1 (brown) /EDTA, (D) TIA1 with ZnCl_2_ /no EDTA, and (E) TIA1 with TC1 and ZnCl_2_ /no EDTA. The brown line represents TC1. In figure E, we assume TC1 cannot bind to RRM3 because of competition with Zn^2+^.

Panel D shows a model of oligomeric Zn^2+^ bound forms of TIA1. TIA1:Zn^2+^ oligomers appear as large irregular droplets with an average dimension of 50 nm (Fig. 3C and 18D). Both RRM2 and RRM3 have multiple predicted partial Zn^2+^ binding sites as suggested in Fig 10. A stable coordination of Zn^2+^ is likely to require participation of two partial binding sites. Presumably, Zn^2+^ connects two partial binding sites from two monomers and builds up an interdomain interaction network between different RRM2s and RRM3s, resulting in rapid oligomerization and intensive turbidity change.

Panel E shows a model of the oligomers of fl TIA1 bound to both TC1 and Zn^2+^. TIA1:TC1:Zn^2+^ oligomers form round particles with a size smaller than TIA1:Zn^2+^ oligomers (Fig. 3D and 18E). In the TIA1:Zn^2+^ complex, Zn^2+^ binds to RRMs from different TIA1 monomers. Possibly, the addition of TC1 to the TIA1:Zn^2+^ complex interrupts the crosslink of two partial Zn^2+^ binding sites. Especially, considering RRM2 has a strong binding affinity to TC1 but not Zn^2+^, TC1 bound to RRM2 is likely to re-organize the partial Zn^2+^ binding site in RRM2 and result in a collapse of the large TIA1:Zn^2+^ oligomers. But the linkage between Zn^2+^ binding with two RRM3 from different monomers might still remain resulting in a TIA1:TC1:Zn^2+^ complex size that is larger than apo TIA1 but smaller than TIA1:Zn^2+^ complex. In this model the metal might act in a regulatory role, selecting for nucleic acids that have affinity for RRM2 but not for RRM3.

## DISCUSSION

The NMR and ITC data presented here clarified that oligomeric full length TIA1 (prepared in vitro) is a functional form, in that it is capable of binding nucleic acids and/or Zn^2+^. Moreover, solid-state NMR data allowed us to identify specific residues involved in binding, and suggested that changes in interdomain packing occur upon binding. ssDNA binding caused residues close to the predicted DNA binding site in RRM2, such as V137, to undergo chemical shift perturbations, confirming that this binding mode is operative in the full-length oligomeric form. Additional shifts upon binding are seen for residues such as V205 and V206 in RRM3, distal to the DNA binding site, suggesting allosteric remodeling effects upon ligation. Addition of Zn^2+^ resulted in shifts in residues close to the predicted in RRM3 Zn^2+^ binding site, as well as distal residues such as S146 and other residues in RRM2, confirming that the predicted sites in RRM3 are the main site for Zn^2+^ binding in the oligomeric full-length form. Given that a full Zn^2+^ binding shell requires at least two predicted Zn^2+^ binding sites, Zn^2+^ may presumably cross-link two RRM3s from two different monomers and induce rapid oligomerization. Alternatively, it could occupy the Zn^2+^ binding site around S146 in RRM2 together with another Zn^2+^ binding site from RRM3.

We show additional evidence from scattering and electron microscopy that binding of key ligands, Zn^2+^ and nucleic acids, modulates the supramolecular structure of TIA. Zn^2+^ accelerates oligomerization and phase separation of TIA1, presumably by reversibly colocalizing metal binding half-sites from various domains (at least in part intermolecularly), resulting in a notable increase in the size of the supramolecular complexes from 20 to 50 nm. Nucleic acids affect oligomerization in a more complex fashion, dependent on the sequence and the presence vs absence of Zn^2+^. ssDNA with a single binding site binds to TIA1 with high affinity, with relatively little effect on oligomerization.

Longer lengths of ssDNA with multiple binding sites interspaced by linkers, on the other hand, “daisy-chain” the TIA1 to form long multi-component strands. The combination of both ligands, though, had an interesting feature. As compared to TIA1:TC1, reversible Zn^2+^ binding to form TIA1:TC1:Zn^2+^ caused the complex to become compact and shield the nucleic acids from solution. Based on the binding motifs evidenced in our NMR data, this is likely because Zn^2+^ blocks DNA from binding to RRM3, and therefore discourages “daisy chain” topologies. The dramatic restructuring of the supramolecular organization of nucleic acids upon Zn^2+^ binding highlights a potentially regulatory role for Zn^2+^ binding to TIA, wherein Zn^2+^ unlocks the potential of TIA1 to remove transcripts and other nucleic acids from the cellular traffic.

Although in this study we investigated full-length TIA1, the NMR data did not display peaks for RRM1 and the low complexity domain regardless of the ligands added. If the protein formed amyloids, strong NMR signals characteristic for beta sheet conformations would have been expected. Together with the EM images the data make a strong case that the wild-type full-length protein does not tend to form amyloid fibrils, even in presence of ligands. There are however two caveats to consider regarding this observation. Truncated forms containing only the LCD are known to form amyloids in vitro, so proteolytic processing or alternative splicing would be expected to lead to nucleation of amyloids. Also, conventional protein-expression systems with *E. coli* cells do not possess specific enzymes involved in eukaryotic post-translational modifications. However, the ability of RNA-binding proteins to assemble into biomolecular condensates can be dramatically affected by post-translational modifications in eukaryotic cells.(16) Post-translational modifications alter protein folding or molecular interactions by changing the charge, hydrophobicity and size of the residues through additions of functional groups (e.g. phosphorylation, acetylation, methylation, etc.) or small protein fraction (e.g. SUMOylation and ubiquitination).(33) Many RNA-binding proteins have been identified that undergo a significant number of post-translational modifications, and some are considered hallmarks of neurodegenerative diseases. Aberrant post-translational modifications lead to loss of physiological function and irreversible phase separation. For example, in the case of TDP-43, phosphorylation of S409 and S410 in the intrinsically disordered domain is a highly consistent abnormal event observed in disease states.(34) Mutation of S409/410 into phosphomimetic aspartic acid residues to mimic the phosphorylation in vitro significantly reduces the oligomerization of TDP-43 by introducing electrostatic repulsion. In FUS-associated frontotemporal lobar degeneration, FUS is hypomethylated and accumulates in neurons. Arginine hypomethylation strongly promotes phase separation of FUS by regulating cation-p interactions between C-terminal arginines and N-terminal tyrosines.(35) Although TIA-1 has also been proved to undergo post-translational modifications during Fas-mediated apoptosis(36), relatively little information is available about the effect of post-translational modifications of TIA1 on stress granule assembly. It would be interesting to probe whether specific TIA1 post-translational modifications events interfere with its cooperative association to mRNAs and then up- or down-regulate oligomerization.

Finally, although wild type TIA does not form amyloid fibrils, neither in the apo nor in the DNA and Zn^2+^ ligated forms, the situation may be different for disease-associated mutations, most of which are found in TIA1 lie in the C-terminal low complexity domain (P362L, A381T, or E384K).(8) These mutations may alter biophysical properties of TIA1, strengthening the intermolecular interactions and dramatically altering the functions in the cell. For example, in some models leading to the impaired disassembly of stress granules, possibly contributing to the formation of irreversible amyloid fibrils. MD simulations support the hypothesis that P362L has a strong tendency to induce β-sheet interactions.(37) Additional work is needed to investigate the molecular mechanisms for how these mutations disturb the dynamics and function of TIA1 in vitro and stress granules in vivo.

## CONCLUSIONS

We previously characterized the reversibly formed oligomeric micelle-like form of full length apo mouse TIA1. Here, we applied solid-state NMR to obtain atomic resolution evidence for structural alterations of the RRM domains in full-length TIA1 upon binding with oligonucleotides. When oligomeric full-length TIA1 was complexed with ssDNA or Zn^2+^ (or both), specific side-chain atoms of TIA1 exhibited NMR chemical shift perturbations. Evidence of DNA binding was observed in RRM2 while evidence of Zn^2+^ binding was observed in RRM3. Notably, we saw no evidence for formation of amyloid fibrils, nor any other dramatic conformation change, regardless of whether Zn^2+^ or DNA was bound. These observations provide clear evidence that the in vitro oligomeric form of fl TIA1 is competent for nucleic acid binding, and that binding may affect oligomerization of TIA1, highlighting the possibility that this form of TIA1 may serve as a good model for its function in stress response and neurodegenerative diseases. Different supramolecular organization was observed depending on the ligands present, as attested by EM data. The NMR and EM suggest a potential regulatory role for Zn^2+^, which induces a compact form of the TIA1:DNA complex, effectively shielding the DNA from solution.

## AVAILABILITY

Raw NMR data files, processing, and analysis scripts are available upon request.

## SUPPLEMENTARY DATA

Supplementary Data are available at NAR online.

## ACKNOWLEDGEMENT

The authors thank Jia Ma, Eric Keeler (NYSBC), Kirk Martin Baughman and Emme Marie Pogue for helpful discussions. The plasmid for the protein expression was provided by the laboratory of Eric Kandel.

## Author contributions

Y.Y., K.J.F. and A.E.M. designed research; Y.Y. and S.H. prepared the NMR samples; Y.Y. and K.J.F. contributed to the NMR data collection and analysis; Y.Y. performed the EM experiments; Y.Y. and S.H. contributed to the oligomerization assay and ITC studies; Y.Y. and A.E.M. wrote the paper; and A.E.M. supervised.

## SUPPLEMENTAL INFORMATION

**Fig S1.**
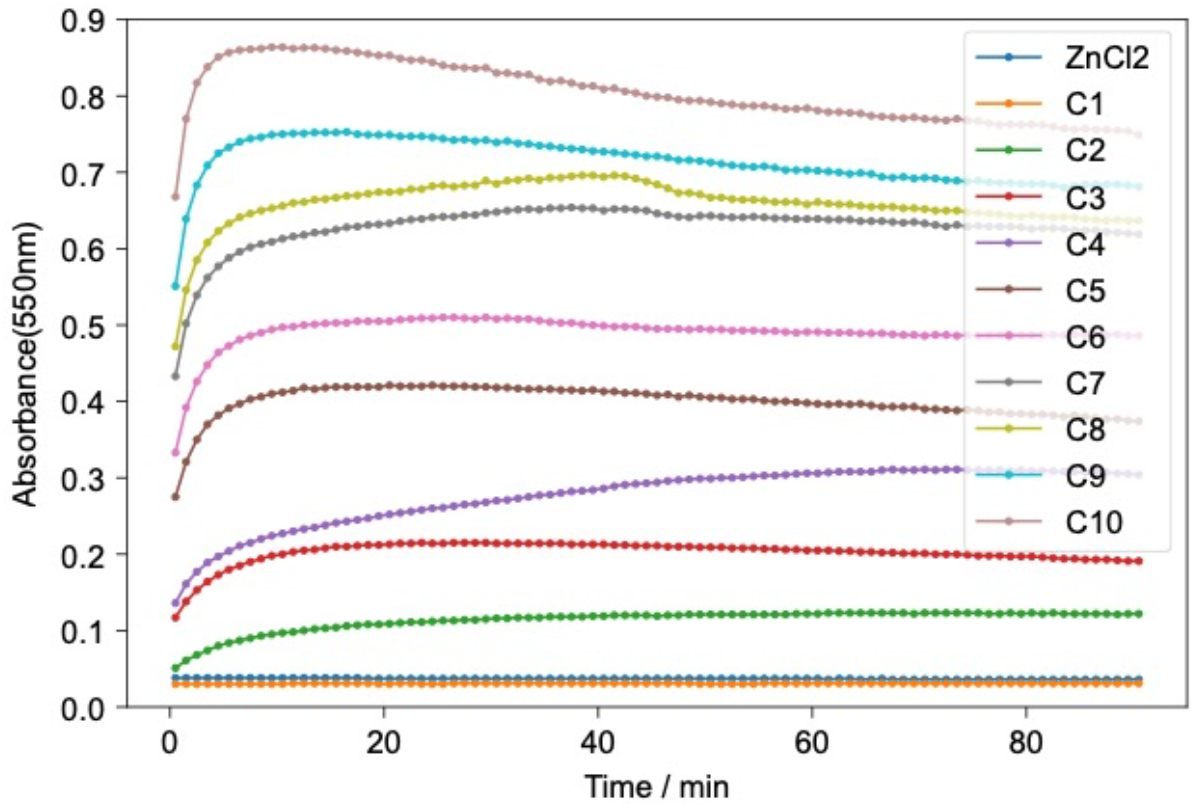
Turbidity (A550 nm) of 0-8 μM TIA1 with excess ZnCl_2_. The exact concentrations of TIA1 in each well are shown in table S1. The concentration of ZnCl_2_ was 350 μM. The buffer contained 25mM PIPES, 50mM NaCl, 5mM TCEP, 0.002% sodium azide, 20% glycerol, pH 6.8. The turbidity of buffer with no protein was measured for reference.

**Fig S2.**
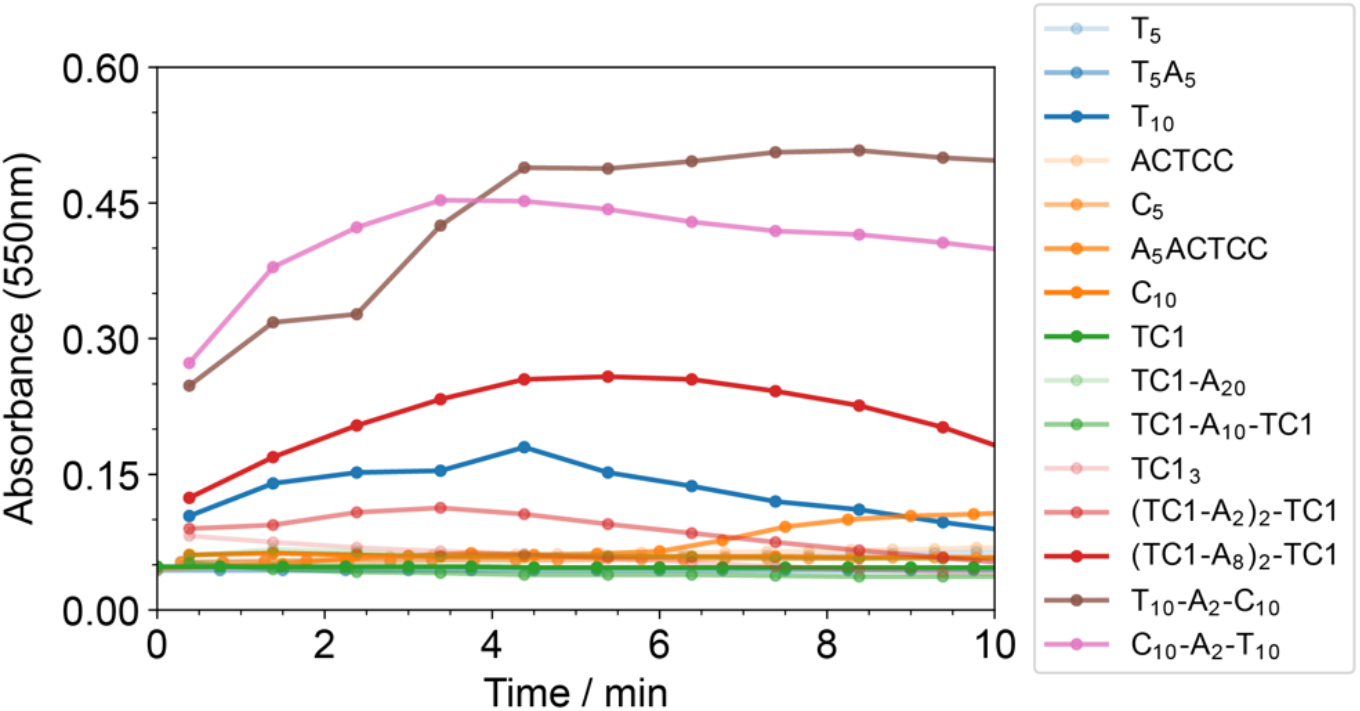
Turbidity of TIA1 with different ssDNA sequences. The DNA sequences used in the oligomerization assay and the maximum turbidity (A550 nm) of each TIA1:DNA complex were listed in table 3. Both protein and ssDNA concentration were at 20 μM. ssDNA with multiple binding sites caused more oligomerization regardless of the length of the nucleotides.

**Table S1.**
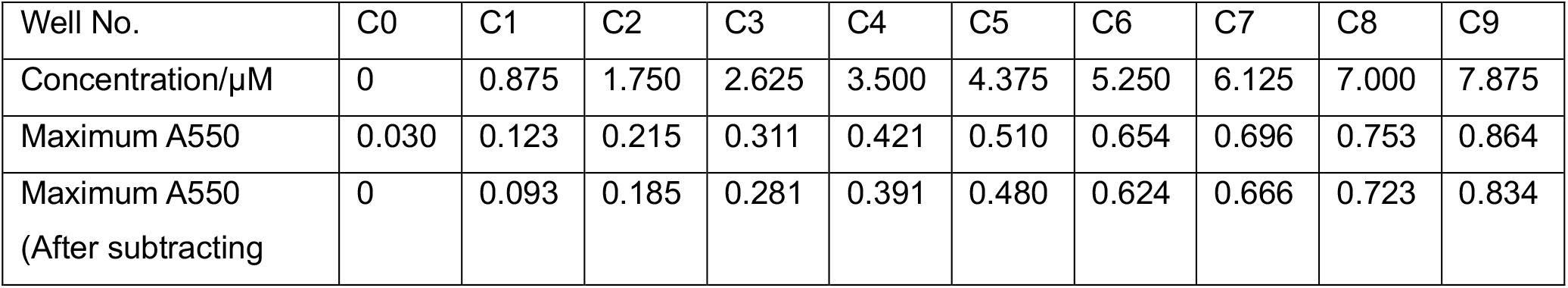
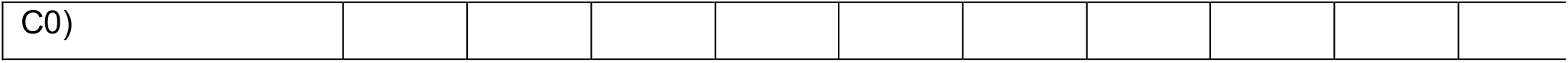
Increasing concentration of TIA1 oligomers in the presence of Zn^2+^

**Table S2.**
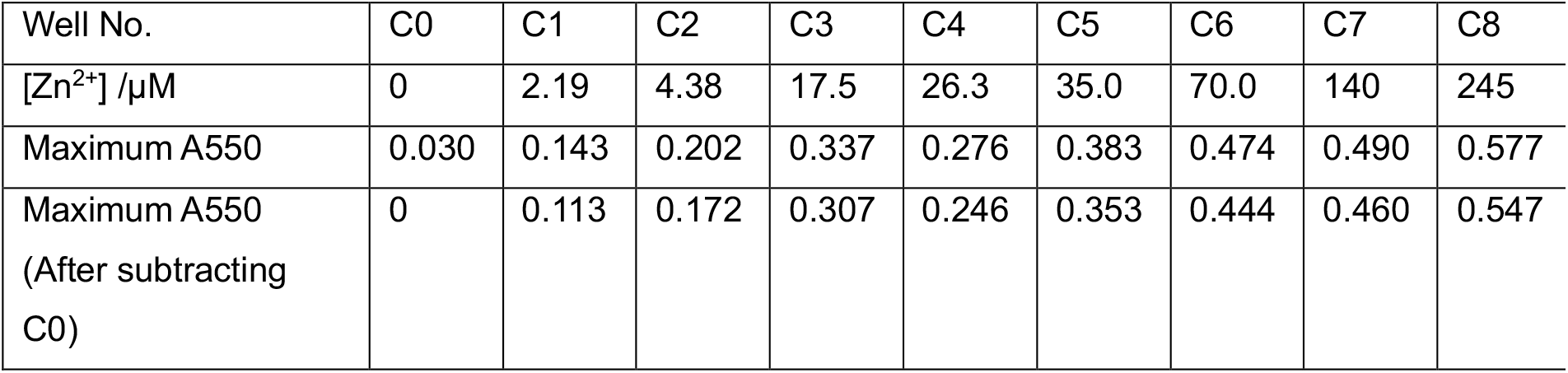
Increasing concentration of TIA1 oligomers in the presence of Zn^2+^

## Sequence alignment

**Fig S3.**
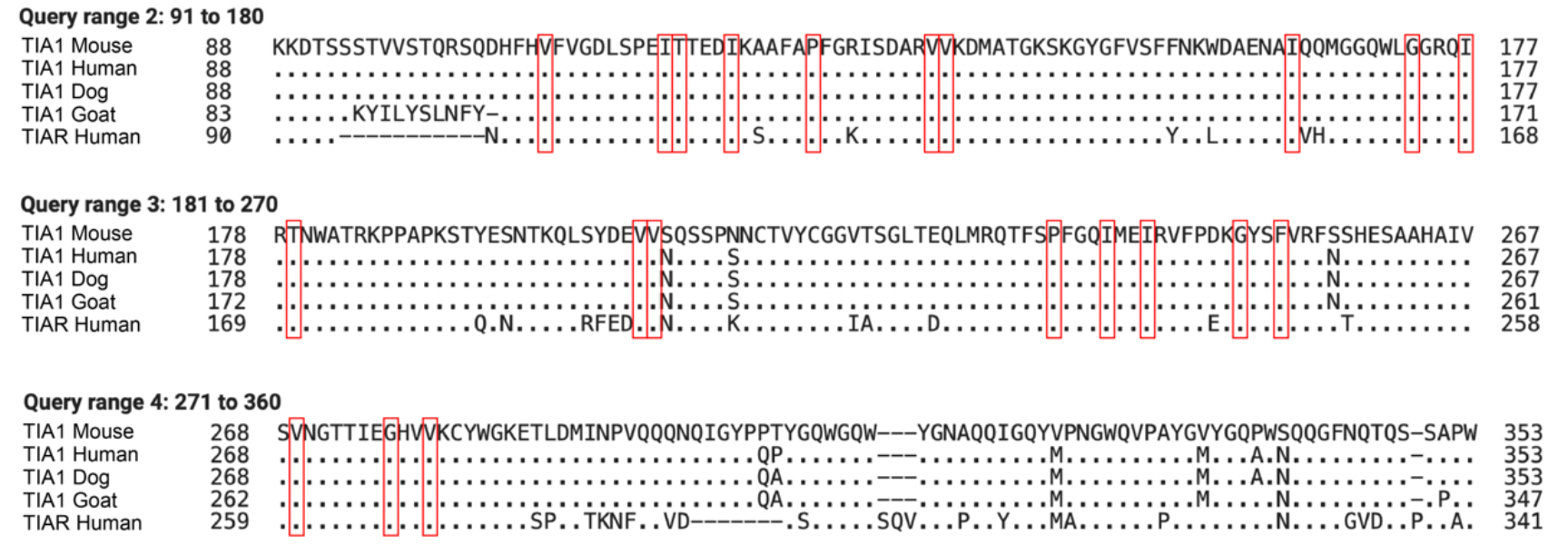
Sequence alignment of TIA1 isoforms. The residues with chemical shifts changes upon binding to TC1 were highlighted with red boxes. They are all strongly conserved.

## Sample preparation

When the concentration of the protein was higher than 1.2 mg/mL, it became opalescent or opaque at room temperature, and this was reversible if the protein was diluted. In figure S4A, after adding Zn^2+^, TIA1 was quickly oligomerized (within 5 min), and the sample was no longer homogenous. The oligomers settled down to the bottom of the tube without spinning. This Zn^2+^-induced TIA1 oligomerization can be partially reversed by adding EDTA. In figure S4B, after adding 10nt TC1 ssDNA into TIA1, the solution was clearer than before. This phenomenon agreed with the results of oligomerization assay as well. In figure S4C, after adding Zn^2+^ to the mixture of TIA1 and TC1, the turbidity increased again, but the not enough to result in particle settle with gravity (in absence of centrifugation). All of samples were stirred for two days at 4°C before packing into an NMR rotor, regardless of whether prompt oligomerization was apparent by eye.

**Fig S4.**
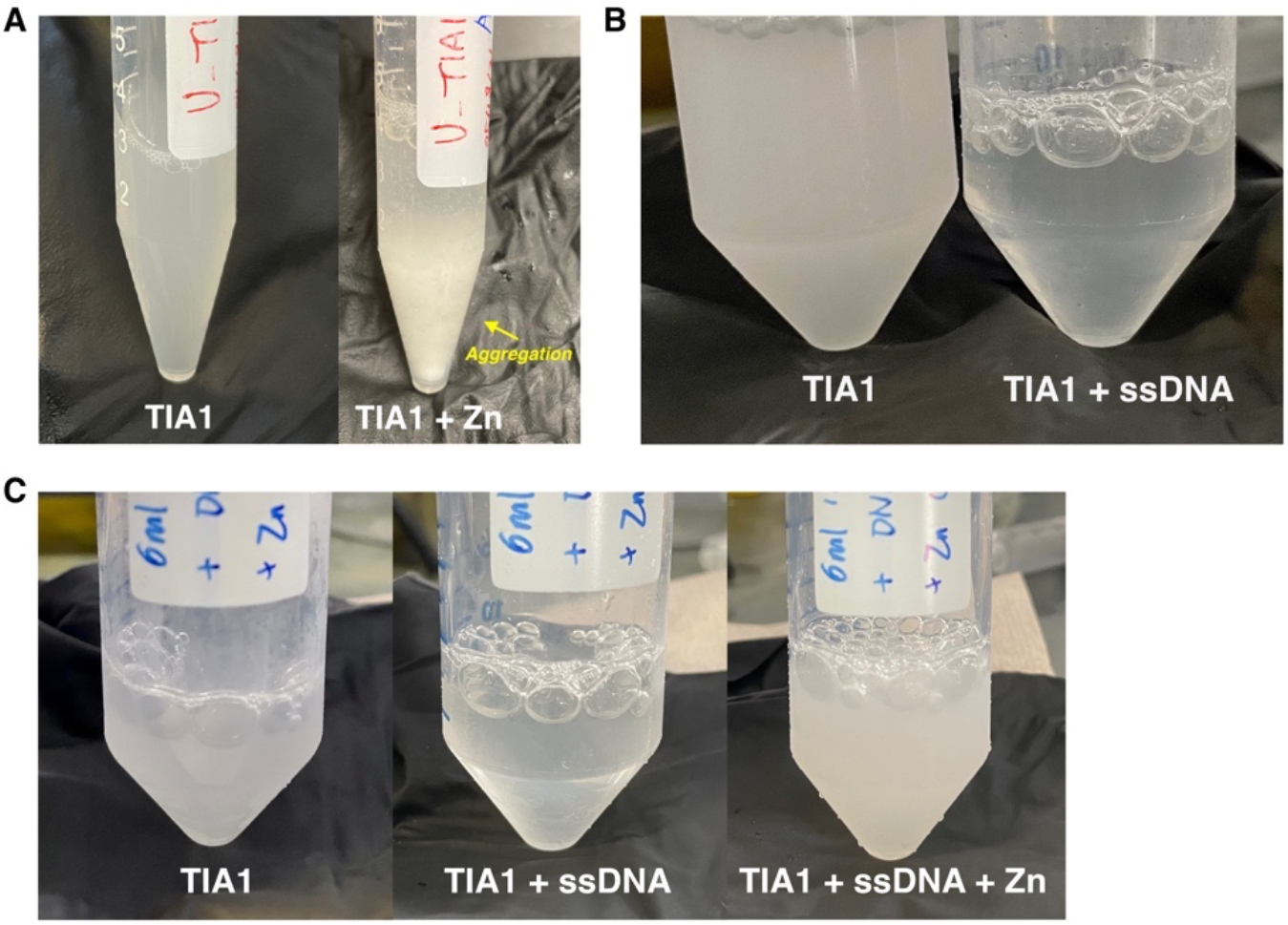
Samples before centrifugation and packing into an NMR rotor. (A) Comparison between TIA1 and TIA1 with Zn^2+^. (B) Comparison between TIA1 and TIA1 with TC1 ssDNA. (C) Comparison among (1) TIA1, (2) TIA1 with TC1 ssDNA, and (3) TIA1 with TC1 ssDNA and Zn^2+^.

