## supplemental figures for "Zinc Alters the Supramolecular Organization of Nucleic Acid Complexes with Full-Length TIA1"

**SUPPLEMENTAL INFORMATION**

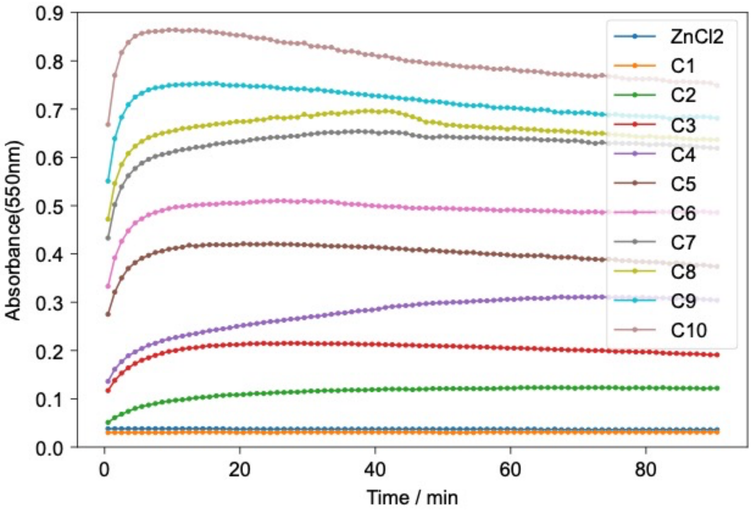

Fig S1. Turbidity (A550 nm) of 0-8 μM TIA1 with excess $ZnCl_2$. The exact concentrations of TIA1 in each well are shown in table S1. The concentration of $ZnCl_2$ was 350 μM. The buffer contained 25mM PIPES, 50mM NaCl, 5mM TCEP, 0.002% sodium azide, 20% glycerol, pH 6.8. The turbidity of buffer with no protein was measured for reference.

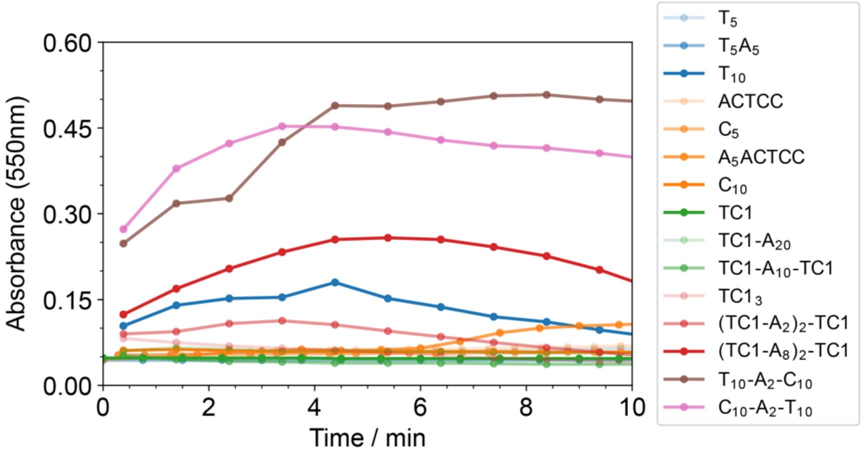

Fig S2. Turbidity of TIA1 with different ssDNA sequences. The DNA sequences used in the oligomerization assay and the maximum turbidity (A550 nm) of each TIA1:DNA complex were listed in table 3. Both protein and ssDNA concentration were at 20 μM. ssDNA with multiple binding sites caused more oligomerization regardless of the length of the nucleotides.

Table S1. Increasing concentration of TIA1 oligomers in the presence of $Zn^{2+}$

| Well No. | C0 | C1 | C2 | C3 | C4 | C5 | C6 | C7 | C8 | C9 |
|---|---|---|---|---|---|---|---|---|---|---|
| Concentration/μM | 0 | 0.875 | 1.750 | 2.625 | 3.500 | 4.375 | 5.250 | 6.125 | 7.000 | 7.875 |
| Maximum A550 | 0.030 | 0.123 | 0.215 | 0.311 | 0.421 | 0.510 | 0.654 | 0.696 | 0.753 | 0.864 |
| Maximum A550<br>(After subtracting | 0 | 0.093 | 0.185 | 0.281 | 0.391 | 0.480 | 0.624 | 0.666 | 0.723 | 0.834 |

| |
|---|
| C0) |

Table S2. Increasing concentration of TIA1 oligomers in the presence of $Zn^{2+}$

| Well No. | C0 | C1 | C2 | C3 | C4 | C5 | C6 | C7 | C8 |
|---|---|---|---|---|---|---|---|---|---|
| $[Zn^{2+}]$ /μM | 0 | 2.19 | 4.38 | 17.5 | 26.3 | 35.0 | 70.0 | 140 | 245 |
| Maximum A550 | 0.030 | 0.143 | 0.202 | 0.337 | 0.276 | 0.383 | 0.474 | 0.490 | 0.577 |
| Maximum A550 (After subtracting C0) | 0 | 0.113 | 0.172 | 0.307 | 0.246 | 0.353 | 0.444 | 0.460 | 0.547 |

### Sequence alignment

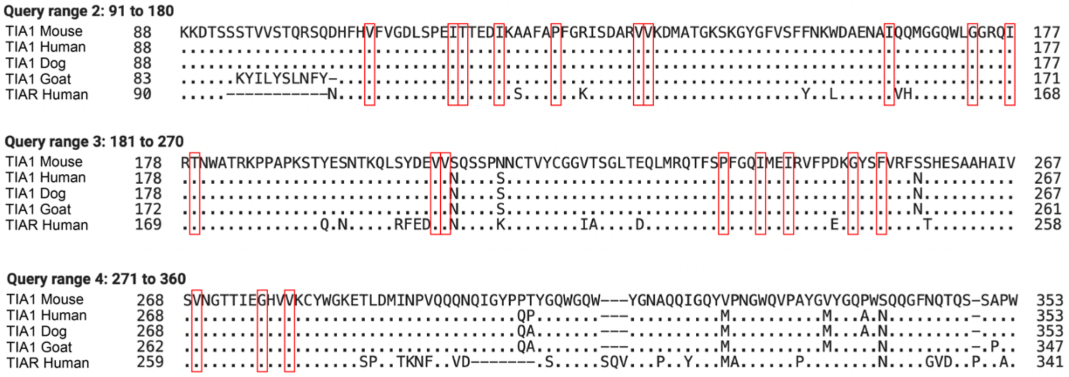

Fig S3. Sequence alignment of TIA1 isoforms. The residues with chemical shifts changes upon binding to TC1 were highlighted with red boxes. They are all strongly conserved.

### Sample preparation

When the concentration of the protein was higher than 1.2 mg/mL, it became opalescent or opaque at room temperature, and this was reversible if the protein was diluted. In figure S4A, after adding $Zn^{2+}$, TIA1 was quickly oligomerized (within 5 min), and the sample was no longer homogenous. The oligomers settled down to the bottom of the tube without spinning. This $Zn^{2+}$-induced TIA1 oligomerization can be partially reversed by adding EDTA. In figure S4B, after adding 10nt TC1 ssDNA into TIA1, the solution was clearer than before. This phenomenon agreed with the results of oligomerization assay as well. In figure S4C, after adding $Zn^{2+}$ to the mixture of TIA1 and TC1, the turbidity increased again, but the not enough to result in particle settle with gravity (in absence of centrifugation). All of samples were stirred for two days at 4°C before packing into an NMR rotor, regardless of whether prompt oligomerization was apparent by eye.

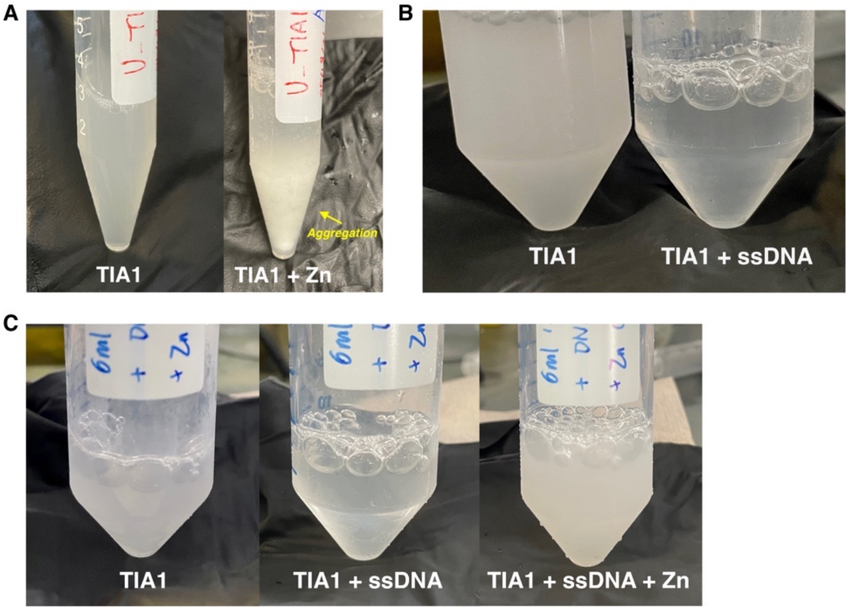

Fig S4. Samples before centrifugation and packing into an NMR rotor. (A) Comparison between TIA1 and TIA1 with $Zn^{2+}$. (B) Comparison between TIA1 and TIA1 with TC1 ssDNA. (C) Comparison among (1) TIA1, (2) TIA1 with TC1 ssDNA, and (3) TIA1 with TC1 ssDNA and $Zn^{2+}$.
